# Exploring brain lobe-specific insights in an explainable framework for EEG-based schizophrenia detection

**DOI:** 10.1101/2025.09.29.679358

**Authors:** Md. Milon Hossain, Md. Nurul Ahad Tawhid

**Author notes:** These authors contributed equally to this work.

## Abstract

Schizophrenia (ScZ) is a growing global health concern that affects millions of people and puts severe pressure on healthcare systems. Early detection and accurate diagnosis are crucial for adequate management. Electroencephalography (EEG) has evolved into a promising non-invasive tool for detecting ScZ in contemporary research. However, specific biomarkers, especially those related to brain lobes, cannot often be identified by current EEG-based diagnostic methods. Different brain lobes are associated with distinct cognitive functions and patterns of diseases. Also, there is a gap in the incorporation of the XAI technique, as medical diagnosis needs trustworthiness and explainability. This study strives to address these gaps by developing a framework using mel-spectrogram images with Convolutional Neural Networks (CNNs). EEG signals are converted into mel-spectrogram images using Short-Time Fourier Transform (STFT). After that, these images are analyzed using a CNN model to perform classification between ScZ and healthy control (HC). To identify the most critical brain regions, the full brain regions are divided into five different regions, and the same classification process is performed. The performance of the proposed framework is evaluated using two publicly available EEG datasets: repOD and the kaggle basic sensory task dataset, which provides a remarkable accuracy of 99.82% and 98.31% respectively. Among regions, the frontal lobe has the most significant performance with an accuracy of 97.02% and 88.03%, respectively, in these datasets, followed by the temporal lobe. Conversely, the occipital lobe shows the lowest accuracy among lobes, with only 79.30 % and 68.33% accuracy on both occasions, showing its lower significance in the diagnosis. To bring result explainability, LIME, SHAP, and the Grad-CAM methods are applied, providing valuable insights for clinicians and researchers. These findings emphasize the potential of EEG-based brain lobe analysis in enhancing ScZ detection, diagnostic accuracy, explainability, and clinical guidance.

## Introduction

Schizophrenia (ScZ) is a chronic, disabling neuropsychiatric disorder characterized by disturbances in thought, perception, emotions, and behavior. It affects nearly 24 million people, or around 0.33% of the global population, according to the latest World Health Organization (WHO) report [1]. Individuals with ScZ often experience persistent delusions, hallucinations, disordered thinking, cognitive problems, poor social functioning, and reduced quality of life, which cause considerable daily challenges [1]. Although SCZ is a complex disease, it is essential to develop an effective treatment system and enhance patient outcomes. A thorough understanding and management of this complex disease requires a holistic approach that integrates biological, psychological, and social factors [2]. A survey indicated that annual costs for the diagnosis of ScZ patients in the country ranged from US $94 million to US $102 billion, and indirect costs contributed to 50%–85% of the total costs associated with this disease [1]. The increasing frequency imposes a considerable strain on healthcare systems and economies. The economic burden of ScZ was estimated to range from 0.02% to 1.65% of the gross domestic product in most high-income countries. Over the years, it has been increasing rapidly [3, 4]. Despite its devastating consequences, ScZ lacks a cure or treatment capable of preventing or stopping its advancement. Nonetheless, early identification can alleviate its consequences and enhance patients’ quality of life. [5].

A few techniques are available for capturing the functional activity of the brain. Conventional diagnostic procedures for ScZ, including clinical evaluations and neuroimaging modalities (MRI, PET scans), are costly, labour-intensive, and not extensively available [6–9]. Consequently, researchers are progressively investigating non-invasive, economical alternatives, such as electroencephalography (EEG) [8–11]. It provides significant benefits due to its superior temporal resolution, cost-effectiveness, user-friendliness, and direct assessment of neural activity [12]. EEG records the brain’s electrical activity using electrodes attached to the scalp [12]. Research has repeatedly demonstrated that persons with ScZ display unique EEG patterns, characterized by modified power spectra, coherence, and connectivity. These correlate with cognitive impairments, sensory processing deficiencies, and other clinical symptoms, especially in several brain lobes [13–15]. The human brain comprises multiple lobes—frontal, temporal, parietal, occipital, and central each essential for cognitive function [13]. Temporal lobe disturbances are frequently linked to memory and language problems in ScZ disease [14], whereas variations in the frontal and parietal lobes may signify cognitive impairment and executive dysfunction [15]. Research has associated EEG abnormalities in particular lobes with neurodegeneration leading to ScZ. There is growing evidence that employing EEG data to identify lobe-specific neural biomarkers could provide an essential aspect in the domain of ScZ diagnosis.

In recent years, numerous studies have concentrated on identifying ScZ by EEG data [16–45]. These studies exploited various machine learning algorithms in conjunction with diverse feature extraction methods for EEG-based ScZ detection. Feature extraction techniques, including time, frequency, and time-frequency analysis [16–18, 21],entropy measures [7],connectivity measures [19],complexity measures [20],and event-related analysis [8]. Then these features have been employed alongside established machine learning models such as support vector machines (SVM) [9, 22–25], k-nearest neighbours (KNN) [4, 26], Boosted trees classifier [27], random forests(RF) [28, 29], PSDL [30, 31], microstate semantic model [32] and Empirical mode decomposition (EMD) [3]. The ML-based classification techniques described above employ a feature engineering approach to create features from raw EEG data, which demands experts with an in-depth knowledge of the feature domain of interest [46]. Conventional ML techniques, with shallow architectures and limited nonlinear feature adaptation, struggle to capture complex patterns in EEG data. To tackle this difficulty, deep learning (DL) approaches have gained prominence because of their capacity to automatically extract and learn features directly from raw EEG data. DL algorithms possess a distinct benefit by automatically executing feature engineering through hidden layers.

The use of DL in EEG data analysis offers potential for precise diagnosis of symptoms and disease advancement. In their 2021 study, Sun et al. [33] used EEG data from 109 subjects (54 with ScZ, 55 with HC) to demonstrate the effectiveness of fuzzy entropy (FuzzyEn) over the Fast Fourier Transform (FFT). It achieved an average accuracy of 99.22% with FuzzyEn and 96.34% with FFT. CNN-based approaches have been widely adopted in DL based methods. The next year, Lillo et al [18] operating through another dataset of 28 subjects (14 with ScZ, 14 HC) achieved 93% accuracy with a CNN model. Using the same dataset, Oh et al. [36] executed an eleven-layer CNN with 98.07% accuracy in non-subject-based and 81.26% in subject-based testing and Latreche et al. [34] reached 99.18% accuracy with a ten-layer CNN. Additionally, Yang et al. [35] integrated both deep CNN and fuzzy logic, resulting in a 99.05% accuracy. Compiling on the same dataset of 28 subjects (14 with ScZ, 14 HC), the RDCGRU model was introduced by Sahu et al. [37], which employed several 1-D Convolutional layers with lengths greater than one. It attained an accuracy of 88.88% with alpha-EEG rhythms. Additionally, Goker et al. [16] presented a 1D-CNN with multitaper techniques, which increased the accuracy to 98.76% in the same dataset. 3D CNN models have also shown the best potential, with Guo et al. [45] achieving 99.46% accuracy using a dataset of 28 subjects (14 with ScZ, 14 HC). It further reveals altered brain connectivity and reductions in entropy in the temporal and frontal lobes.

Some studies have transformed EEG into time–frequency images for classification. Using another dataset of 81 samples (49 ScZ and 32 HC), Khare et al. [38] introduced the Continuous Wavelet Transform (CWT), Short-Time Fourier Transform (STFT), and Smoothed Pseudo-Wigner-Ville Distribution (SPWVD) techniques to produce scalograms, spectrograms, and SPWVD-based time-frequency representation (TFR) plots, respectively. The findings demonstrate an accuracy of 93.36% using the SPWVD-based TFR and CNN models. Using the same dataset, Sahu et al. [43] designed a separable convolution attention network (SCZSCAN), which achieved 95% accuracy. Similarly, Aslan et al. [39] utilized STFT-based 2D time-frequency (T-F) images in a DL-based VGG-16 CNN model, which achieved accuracies of 95% and 97.4%, respectively, using two distinct EEG datasets, one of which included 84 subjects (39 HC and 45 ScZ) and the other of which included 28 subjects (14 HC and 14 ScZ). In another study, the same author [40] introduced a method that utilizes the CWT to generate 2D T-F scalogram images. The same VGG16 CNN model was used, achieving accuracies of 98% and 99.5% on previously used datasets. Moreover, techniques such as Activation Mapping, Saliency Map, and Grad-CAM are employed to visualize learning results. Beyond this, using a dataset of 84 individuals (45 ScZ, 39 HC), Naira et al. [41] employed the Pearson correlation coefficient (PCC) with CNNs, achieving an accuracy of 90%. Li et al. [42] applied a lightweight Vision Transformer model (LeViT) with CNN, which obtained a subject-independent performance of 98.99%.

Although some have reached various levels of significance in some cases, their ability for robustness and generalization is limited. None of the previous approaches considered brain lobe-specific diagnosis of ScZ, as different brain lobes are responsible for various functionalities in the human brain. According to recent studies, recognizing lobe-specific EEG biomarkers can enhance the improvement of the diagnosis process [13–15].

In medical decision-making systems, explainable artificial intelligence (XAI) has become an essential element for diagnosing neuropsychiatric disorders, as medical diagnoses require trustworthiness and explainability [47]. Deep learning models offer robust tools for automated classification, but their “black-box” nature limits their interpretability and clinical trust. XAI can bring transparency by highlighting relevant abnormalities that impact model predictions, thereby helping to build reliable diagnostic tools and increasing clinician confidence [48]. Our goal in incorporating XAI into deep learning models is to identify the reasons behind classification decisions. Most importantly, focusing on lobe-specific EEG modifications, we aim to enhance diagnostic precision and aid in the development of non-invasive, accessible, and reliable instruments for the early detection of ScZ.

In this study, to address the existing research gap, we have employed a framework for detecting ScZ and evaluated which brain region is most influential in diagnosis. First, by performing some preprocessing steps using STFT, the EEG signals were converted into mel spectrogram. Then, the converted mel-spectrogram images are fed into CNN models with 10-fold cross-validation. Following the same methodology, we classified the mel spectrograms from each brain lobe and the entire EEG channel set independently to assess how well each approach worked for identifying ScZ. The results obtained from the proposed methods are compared with those from other existing literature that used the same dataset. Notably, three XAI techniques are implemented to introduce explainability to the proposed framework and its outcomes. This research contributes to these noteworthy contributions as follows:

- **Novel biomaker development:** It introduces a novel framework for ScZ detection that integrates mel-spectrogram base images with CNN model, capable of distinguishing between ScZ vs HC using EEG data.
- **Innovative approach:** This research represents the first application of STFT-based mel-spectrogram images in conjunction with a CNN model for the detection of ScZ.
- **Analysis of brain regions:** It explores which brain regions are essential for extracting representative information for accurate ScZ detection.
- **Improvement of performance:** The study seeks to improve the effectiveness of ScZ detection relative to current methodologies.
- **XAI integration:** The study utilizes SHAP, LIME, Grad-CAM to explain the model’s decision-making process, highlighting why different brain regions perform differently in ScZ detection

The following sections of this study are organized as follows: the “Methodology” section outlines the studied data and the proposed method. The “Experiments and Results” part describes the experimental setup and outcomes, whereas the “Discussion” sub-section analyzes the experimental findings and overall analysis. The “Conclusions” section outlines the paper and addresses future study scope.

## Methodology

This study presents a framework for identifying significant brain lobes associated with the detection of ScZ, utilizing EEG brain signal data. The article introduces an approach that integrates STFT-based mel-spectrogram and CNN to identify critical brain lobes as biomarkers. The system initiates with the preprocessing of EEG signals, applying a butterworth band-pass filter to reduce noise. The EEG data are subsequently divided into smaller time segments. Afterwards, the EEG channels are divided into five brain lobes according to biological principles. Mel-spectrogram images are produced for each brain lobe and a full range of EEG channels with STFT, which offers a time-frequency representation of cerebral activity. The mel-spectrograms are then input into a DL-based CNN to classify ScZ vs HC. The classification is conducted independently on the mel-spectrogram images from each brain lobe and the full channel group. Details of these steps are discussed below:

### Proposed Framework

The integration of STFT based mel-spectrogram images with CNN is highly effective for ScZ detection, as STFT encompasses both the time and frequency domains of the brain’s electrical activity, providing valuable insights into cognitive decline. CNNs are recognized for their ability to identify patterns in 2D data such as mel-spectrograms, excel at detecting complex and sophisticated features related to ScZ. This strategy aims to more effectively detect ScZ-related changes in brain wave patterns. A summary of the proposed framework is shown in Fig. 1, with comprehensive details available in the following subsections.

**Fig 1.**
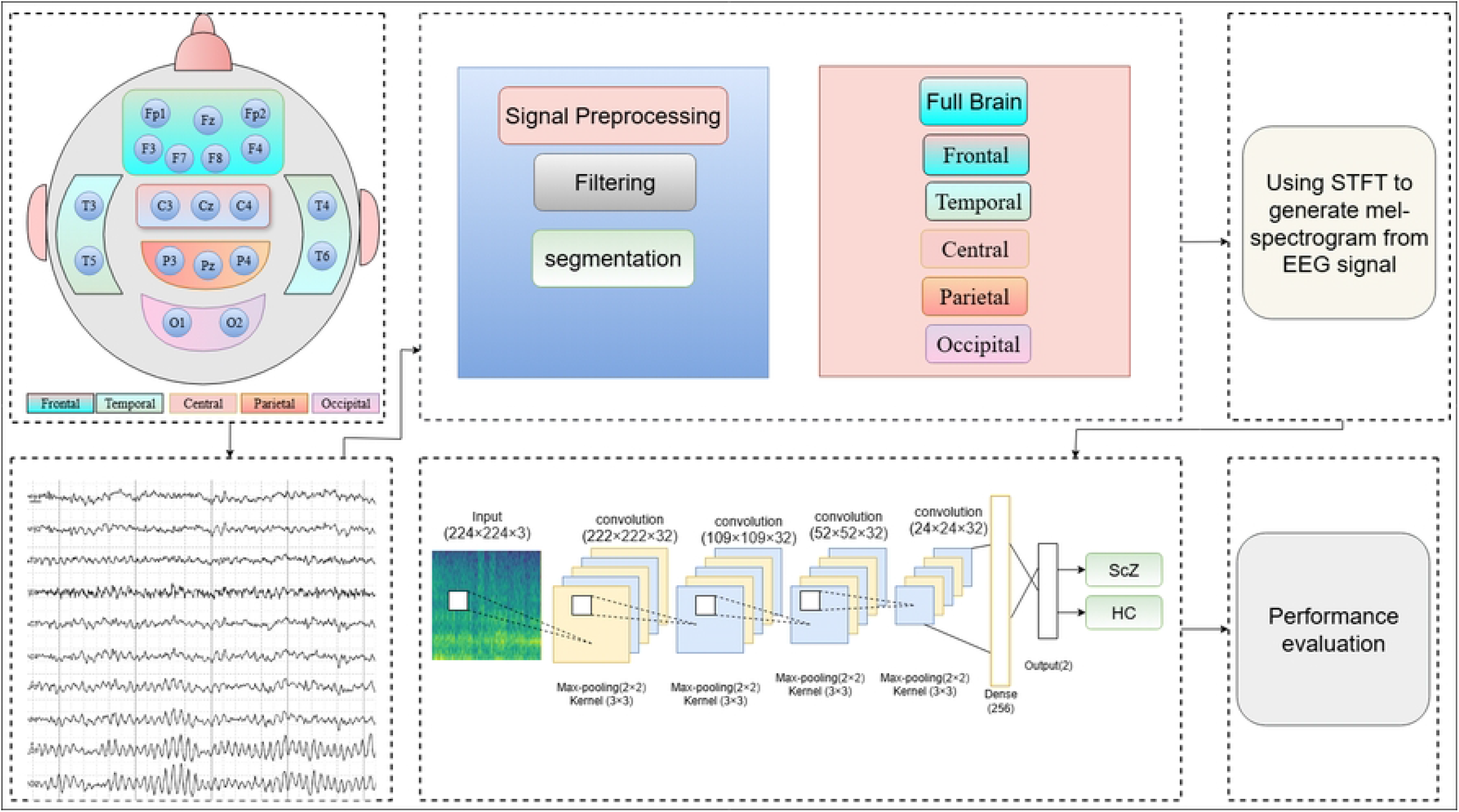
Overview of the proposed framework for detecting ScZ using EEG brain data.

### Dataset description

In our study, two publicly available EEG datasets are used for testing. The first group of data is publicly available called as the repOD dataset and was gathered from the Institute of Psychiatry and Neurology in Warsaw [19]. This dataset includes 19-channel resting state EEGs collected from 28 volunteers of similar ages and genders, half of whom are healthy controls (HC=14) and the other Half (ScZ=14) showed signs of ScZ. The Second dataset, which is comparatively underexplored, was obtained as part of a project financed by the National Institute of Mental Health (NIMH; R01MH058262). This dataset is available to the public on the Kaggle platform [49]. The dataset consists of 64-channel scalp EEG data gathered from 81 Subjects, composed of 49 with ScZ and 32 HCs. A summary of these two datasets is provided in the table 1.

**Table 1.**
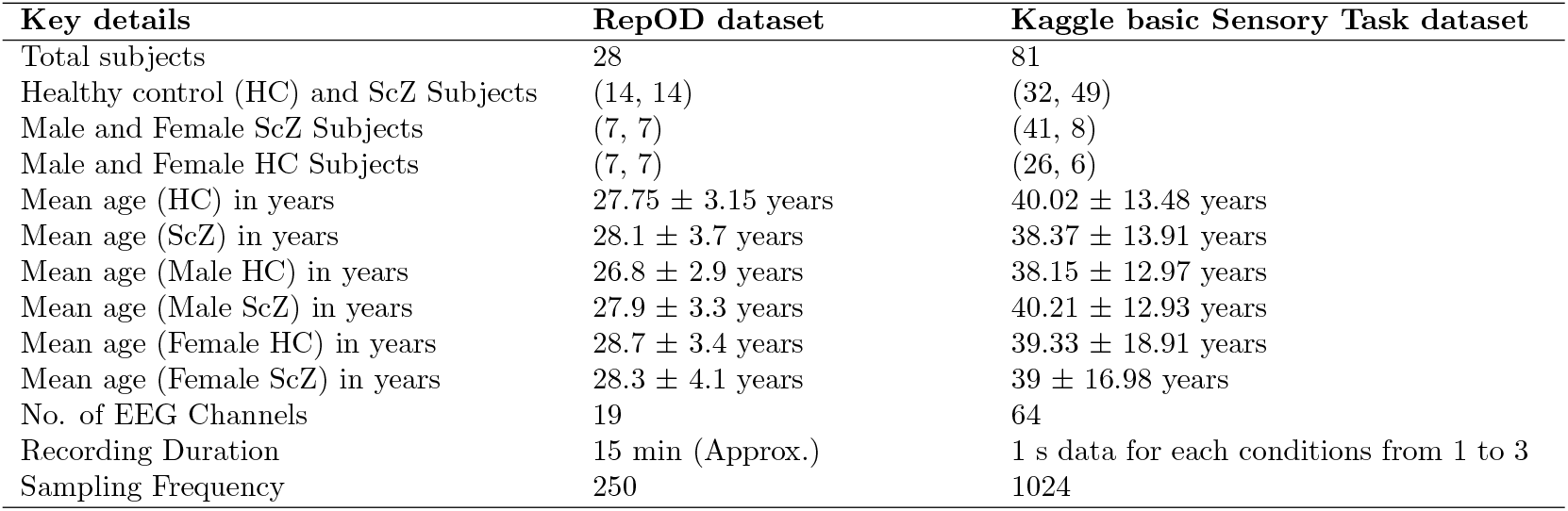
Details of publicly available EEG Schizophrenia dataset.

In the repOD dataset, all subjects were placed in an eyes-closed resting state condition, and their EEG data were captured for 15 minutes. The conventional 10–20 EEG setup with 19 EEG channels, including Fp1, Fp2, F7, F3, Fz, F4, F8, T3, C3, Cz, C4, T4, T5, P3, Pz, P4, T6, O1, O2, was used to collect the data at a sampling frequency of 250 Hz. FCz was the location of the reference electrode. The basic sensory dataset consists of 64 channels, with each file containing 74 columns. The first four columns describe the data, including subject, trial, sample, and condition. The last six columns represent reference/EOG electrodes, such as VEOa, VEOb, HEOL, HEOR, Nose, and TP10. The 64 EEG channels are as follows: ‘Fp1’, ‘Fpz’, ‘Fp2’, ‘AF7’, ‘AF3’, ‘AFz’, ‘AF4’, ‘AF8’, ‘F7’, ‘F5’, ‘F3’, ‘F1’, ‘Fz’, ‘F2’, ‘F4’, ‘F6’, ‘F8’, ‘FC5’, ‘FC3’, ‘FC1’, ‘FCz’, ‘FC2’, ‘FC4’, ‘FC6’, ‘C5’, ‘C3’, ‘C1’, ‘Cz’, ‘C2’, ‘C4’, ‘C6’, ‘FT7’, ‘T7’, ‘TP7’, ‘FT8’, ‘T8’, ‘TP8’, ‘P7’, ‘P8’, ‘P9’, ‘P10’, ‘CP5’, ‘CP3’, ‘CP1’, ‘CPz’, ‘CP2’, ‘CP4’, ‘CP6’, ‘P5’, ‘P3’, ‘P1’, ‘Pz’, ‘P2’, ‘P4’, ‘P6’, ‘O1’, ‘Oz’, ‘O2’, ‘Iz’, ‘PO7’, ‘PO3’, ‘POz’, ‘PO4’, ‘PO8’.

Participants’ anonymity of both datasets are ensured by withholding any personally identifiable information from publication.

### Pre-processing EEG Signals

- **Noise removing:** In repOD, a butterworth filter of order 2 was implemented to filter each EEG channel’s signals in the physiological frequency bands listed below: Delta is 2–4 Hz, theta is 4.5–7.5 Hz, alpha is 8–12.5 Hz, beta is 13–30 Hz, and gamma is 30–45 Hz. In kaggle dataset, EEG data underwent preprocessing with a 0.1 Hz high-pass filter, outlier interpolation for noise reduction, re-referencing to the average of the ear lobes, Single-trial epochs were separated, and baseline correction was implemented. Outliers were eliminated, components were extracted by independent component analysis, and noise was mitigated using canonical correlation analysis. These measures guaranteed data that is clean and prepared for analysis. Then, for these two datasets, a butterworth band-pass filter ranging from 0.5 to 45 Hz was applied to enhance signal quality according to the prior research [9, 31].
- **Resampling:** The signals were resampled to 256 Hz to normalize the data, ensuring consistency in the sampling rate across all channels and enabling accurate comparison and analysis between these two datasets. It is the standard practice in prior research, as sampling is usually done in 2^*n*^ times [24, 38].
- **Segmentation:** In our investigation, the signals were eventually segmented to increase the dataset size by dividing the signal data into equal time intervals and each segment is assigned the same label as the original sample [6, 39]. EEG data were segmented into 3-second segments, as is a standard practice in earlier studies [6, 50–52]. These small portions help capture essential patterns in brain activity without introducing excessive amounts of irrelevant information. This enables the model to extract important features from the signal. Additionally, shorter segments speed up processing and reduce the computational load by decreasing the amount of data that computer must process simultaneously, while preserving essential information.

### Brain Lobe Schedule

To explore the significance of each brain region, we organize the EEG channels into the various brain lobes. The electrodes are positioned according to the 10-20 system, a standardized approach that assures accurate and uniform placement on the scalp based on anatomical landmarks, as shown in Fig. 2. The frontal region, linked to higher thinking and decision-making, is represented by Fp1, Fp2, F3, F4, F7, F8, and Fz. The temporal region, associated with memory and auditory processing, included T3, T4, T5, and T6. The occipital region, responsible for visual processing, used O1 and O2. The central region, involved in motor control and sensory integration, is represented by C3, C4, and Cz. The parietal region, related to spatial orientation, included P3, P4, and Pz.

**Fig 2.**
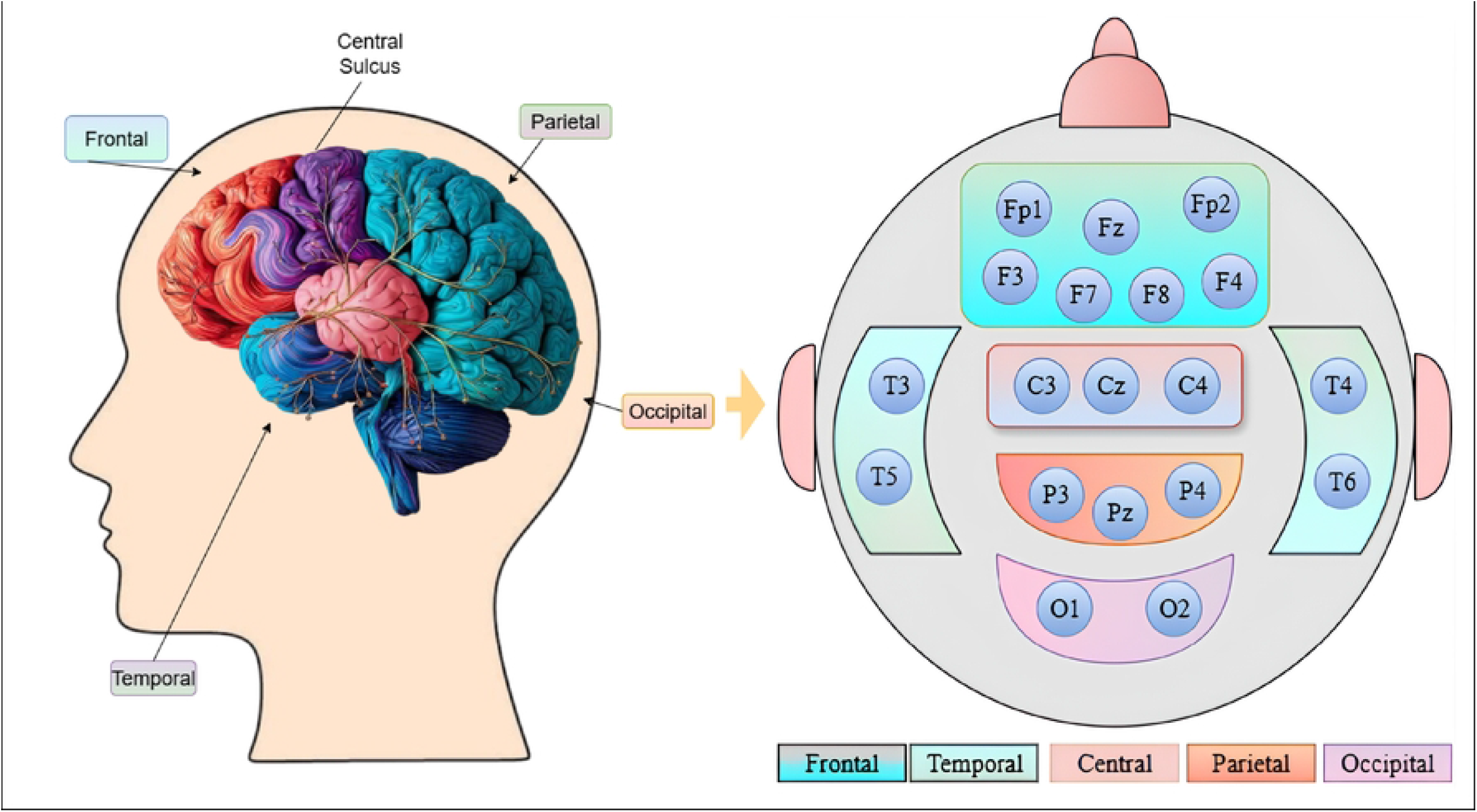
An illustration of the brain lobe structure showing the arrangement of EEG channels for repOD Dataset.

**Fig 3.**
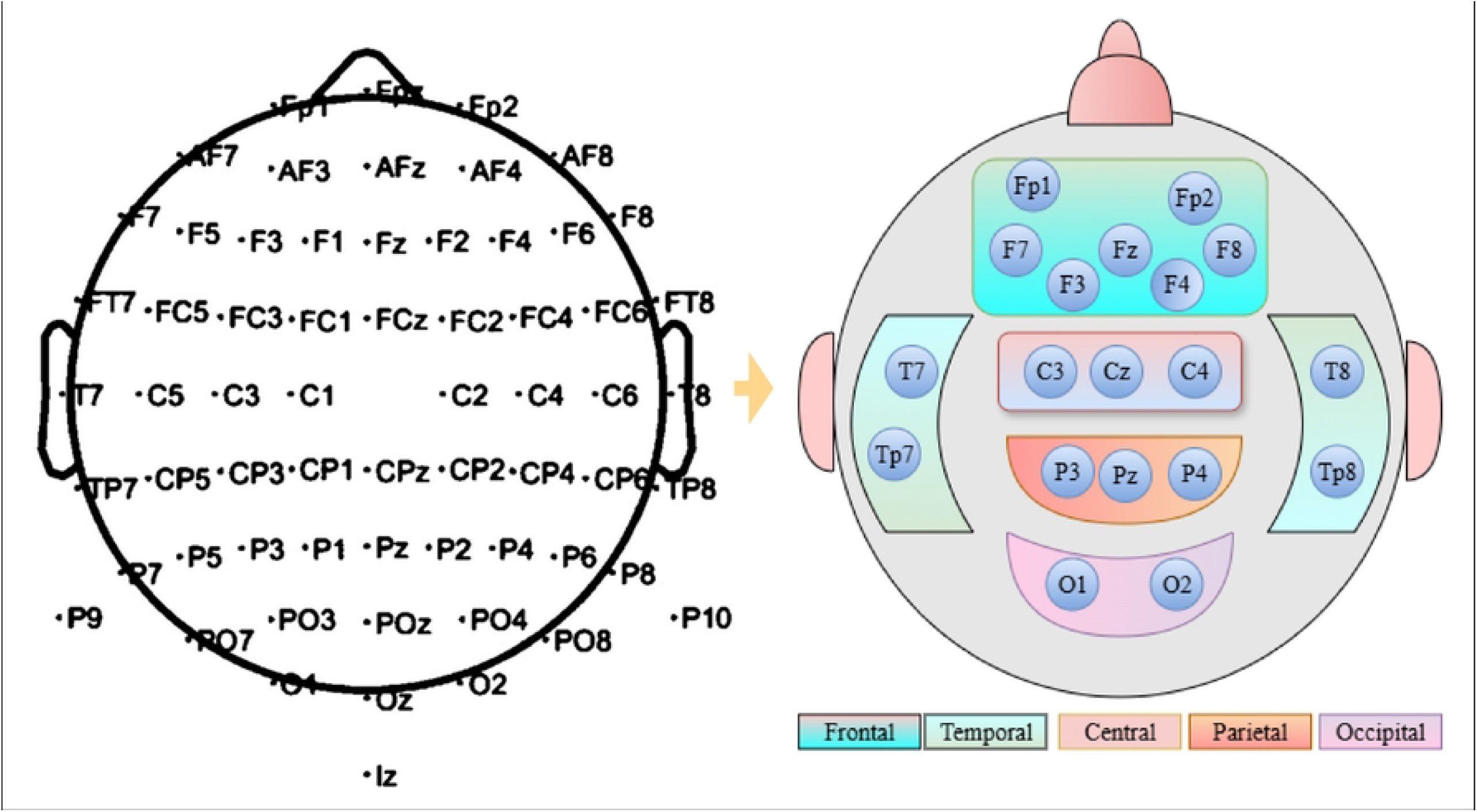
An illustration of the brain lobe structure showing the arrangement of EEG channels for kaggle basic sensory dataset from 64 channels to 19 channels.

The repOD dataset consists of 19-channel electrodes. For the Kaggle dataset, to maintain consistency and clarity, we selected an identical position for 19 channels out of the 64 channels to ensure transparency, as shown in Fig. 4 Then, we divided it into five regions. For the frontal region, the following channels are selected: Fp1, Fp2, F7, F3, Fz, F4, F8. Then, the central region is represented by channels ‘C3’, ‘Cz’, and ‘C4’.

**Fig 4.**
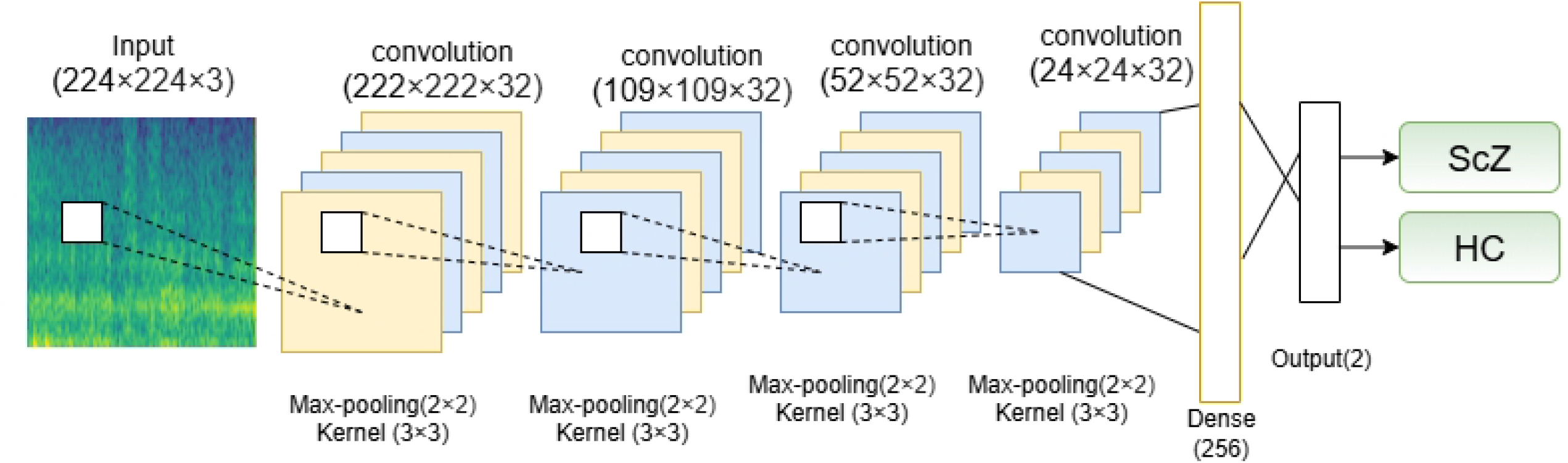
Architecture of the CNN model used in this study.

The temporal and parietal regions are represented by: T7, T8, Tp7, Tp8, and P3, Pz, P4, respectively. Lastly, the occipital region is followed by the channels O1 and O2. Then, all 19 channels are examined as a single set to cover the full region. By organizing the channels based on brain regions, the focus is more precise on how each area contributes to the signals.

### Converting EEG Signals to mel-spectrogram Images

A Mel-spectrogram is a spectrogram in which the frequencies are transformed to the mel scale [5]. The Mel-spectrum comprises an STFT for each spectrum frame (energy/amplitude spectrum), changing from a linear frequency scale to a logarithmic Mel scale, afterwards processed through a filter bank to obtain the eigenvector; these eigenvalues can be approximately represented as the distribution of signal energy across the Mel-scale frequencies. To achieve this, it is initially necessary to calculate the STFT of the signal to convert it into mel-spectrogram images [5, 53]. The STFT is an effective signal processing method used for analyzing the frequency elements in non-stationary signals over time [6]. Following that, the Fourier transform is carried out on each segmented window, producing its specific frequency spectrum. The procedure involves converting the time-varying EEG signal into a two-dimensional (2D) matrix, where time is represented on the horizontal axis and frequency is represented on the vertical axis. The horizontal axis represents time, with the EEG signal divided into segments or windows. The vertical axis denotes the frequency range, generally starting with lower frequencies at the bottom and progressing to higher frequencies at the top [6]. A Hamming window is used for each segment to maintain continuity and minimize spectral leakage. The STFT of a signal at a specific time t and frequency f, denoted as STFT(t, f), is calculated using the following equation:

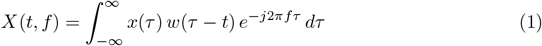

The next phase involves applying a mel filter bank to the STFT output. The mel scale is a logarithmic scale that reflects frequency perceptions, with increased sensitivity to lower frequencies. The mel scale is defined by the following equation:

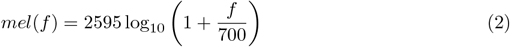

A mel filter bank has overlapping triangular filters that align with various mel frequencies. The filters are placed closely for lower frequencies and more widely for higher frequencies, matching the mel scale [5]. The logarithmic transformation compresses the amplitude range and highlights the more important features of the signal. The outcome is a mel-spectrogram image, a time-frequency representation in which the frequency axis has been converted from linear to mel frequency [53]. The mel-spectrogram illustrates the spectrum of a signal’s energy over time and frequency [5, 53].

### Classification of Schizophrenia: Training the CNN Model with Spectrogram Images

For the classification using mel-spectrogram images, we used a established CNN model from prior research [6]. CNNs are well-known in deep learning for their exceptional efficiency in the execution of classification tasks [16, 34, 45, 50]. They excel in this field by independently finding and extracting relevant features from the input images, thus facilitating accurate classification into several categories [50, 54]. Here is the used CNN model architecture in figure 4.

CNNs accumulate features gradually, beginning with basic patterns and advancing to complex ones. This layered learning process enhances their ability to understand and classify complex visual information with greater precision. When utilized on spectrogram images, adding Mel-mins, which visually represent EEG signals, CNNs can accurately identify and classify them into appropriate groups. Due to their ability to identify complex patterns, CNNs are highly effective and advantageous for numerous image classification tasks [50, 55].

CNNs achieve their efficiency through the specific configuration of their layered architecture. Each convolutional layer contributes individually to feature extraction, enabling the network to learn visual representations hierarchically [6]. Primarily, the layers identify low-level features, including edges, corners, and basic textures. While these features may seem insignificant individually, they constitute the essential components for complex representations. As the data advances through successive layers, the network progressively synthesizes these basic elements into more abstract and semantically significant features [51, 54]. Combinations of edges and textures may create simple geometric shapes that can then become elements in larger objects. As a result, deeper layers in the network can identify high-level patterns and abstract concepts essential to differentiating objects in an image, which enables CNNs to surpass the perception of visual data as merely a structured collection of pixels. The network promotes a systematic and hierarchical approach to interpreting imagery, as it proficiently encodes both local details and global representations of visual content [50, 54].

The convolution process is fundamental in CNNs, as it is essential for extracting significant features from images, thereby facilitating tasks such as image classification and object detection. In a CNN, the input data are processed using 2-D kernels, where convolution is performed by sliding the kernels over the spatial dimensions and computing the sum of dot products between the kernel and the input [6]. The resulting feature maps are then passed through an activation function to introduce non-linearity. The activation value at spatial position (*x, y*) in the *j*^th^ feature map of the *i*^th^ layer, denoted as 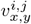, is computed as follows:

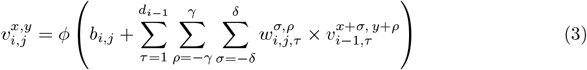

Here, *ϕ* denotes the activation function, and *b*_*i,j*_ is the bias term for the *j*^th^ feature map in the *i*^th^ layer. The term *d*_*i−*1_ represents the number of feature maps in the (*i −* 1)^th^ layer, which also corresponds to the depth of the kernel *w*_*i,j*_. The kernel dimensions are defined by (2*γ* + 1) *×* (2*δ* + 1), where 2*γ* + 1 is its width and 2*δ* + 1 is its height. Finally, 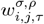 represents the weight parameter of the kernel for the *j*^th^ feature map in the *i*^th^ layer, while 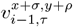 refers to the activation value from the previous layer.

This equation executes a convolution operation by computing the dot product of the filter weights and a localized portion of the input feature map. Integrating the bias, the activation function is employed to produce the final output. The used CNN model aims to effectively extract and learn significant features from input images using a sequence of convolutional and pooling layers. Each convolutional layer uses 32 filters of dimensions 3×3, facilitating the network’s ability to recognize complex spatial patterns. In contrast, max pooling layers with a 2×2 filter are implemented to systematically reduce spatial dimensions and computational complexity without sacrificing critical information. Dropout layers with a rate of 25% are intentionally implemented after the second and fourth convolution-pooling pairs to mitigate overfitting and improve model stability. The combination of convolutional, pooling, and dropout layers assures effective feature extraction while preserving the model’s generalization capability [6].

Then, the network advances to a fully connected (dense) layer including 256 units to combine the acquired features into enhanced representations. A dropout rate of 50% is implemented prior to the final output layer to mitigate overfitting. The classification uses a softmax activation function, generating probabilities for the two target groups: ScZ vs Hc. The model employs categorical cross-entropy loss for training and utilizes the Adam optimizer, resulting in consistent and efficient learning outcomes. This architecture provides a balanced approach to feature learning, regularization, and classification, making it highly suitable for addressing the defined issue.

For a detailed breakdown of the layer configurations, Table 2 illustrates it in a sophisticated way.

**Table 2.** The configuration of the CNN model used in this study.

| Layer type | Filters/units | Kernel size | Strides | Activation | Padding | Dropout rate |
| --- | --- | --- | --- | --- | --- | --- |
| Input | - | - | - | - | - | - |
| Conv2D | 32 | $3 \times 3$ | $1 \times 1$ | ReLU | Same | - |
| MaxPooling2D | - | $2 \times 2$ | $2 \times 2$ | - | - | - |
| Conv2D | 32 | $3 \times 3$ | $1 \times 1$ | ReLU | Same | - |
| MaxPooling2D | - | $2 \times 2$ | $2 \times 2$ | - | - | - |
| Dropout | - | - | - | - | - | 0.25 |
| Conv2D | 32 | $3 \times 3$ | $1 \times 1$ | ReLU | Same | - |
| MaxPooling2D | - | $2 \times 2$ | $2 \times 2$ | - | - | - |
| Conv2D | 32 | $3 \times 3$ | $1 \times 1$ | ReLU | Same | - |
| MaxPooling2D | - | $2 \times 2$ | $2 \times 2$ | - | - | - |
| Dropout | - | - | - | - | - | 0.25 |
| Flatten | - | - | - | - | - | - |
| Dense | 256 | - | - | ReLU | - | - |
| Dropout | - | - | - | - | - | 0.50 |
| Dense(Output) | 2 | - | - | Softmax | - | - |

### Performance Evaluation Methods and Metrics

The main objective of our proposed framework is to accurately assess whether a generated image reflects a HC or one with ScZ, and to figure out which brain region most significantly influences the classification process. For doing so, this research empirically assesses the validation performance through a CNN model. This method facilitates obtaining a trustworthy performance assessment while reducing the possibility of overfitting or underfitting. In standard cross-validation(CV), the training and testing sets vary in each iteration, ensuring that every data point is evaluated. To ensure thorough evaluation and enhanced accuracy, we have opted for a 10-fold CV for our proposed framework. Then, Normalization is introduced to ensure that all input attributes, specifically the pixel values of images, fall within an equivalent range. In this process, each pixel value is divided by 255, resulting in a new range between 0 and 1. It helps in reducing the chances of issues related to differences in scale among input features, ensuring that all data points are treated equally during the model’s training.

To evaluate the model, a series of performance tests is conducted, and the results are analyzed. To thoroughly investigate the efficiency of the proposed solution and methodology, we quantified six established metrics: specificity (Spec), sensitivity (Sen), precision (Prec), accuracy (Acc), F1 score (F1), and false positive rate (FPR). Furthermore, a receiver operating characteristic (ROC) curve serves as a graphical equipment to evaluate the diagnostic performance of the binary classifier system. All metrics are calculated using already established formulas [6, 24]. A ROC curve is a visual representation that is operated to assess the diagnostic accuracy of a binary classification system. It graphs the true positive rate (sensitivity) vs the false positive rate over several threshold settings. The ROC curve is a valuable tool in medical research and machine learning to evaluate a model’s ability to distinguish between two categories, such as the presence or absence of a disease. This enables it to be a crucial tool for assessing predicted precision in various fields, particularly in diagnostic tests and classification challenges [6, 54].

## Experiments and Results

This section begins with a description of the experimental setup, followed by a thorough discussion of the results. Then, it concludes with a complete study of the results analysis.

### Experimental Setup

In the experiment, we used two datasets, both containing two categories of subjects, and carried out classification tasks with ScZ and HC. The basic sensory dataset consisted of 49 patients with ScZ and 32 with HCs. We have used a segmentation length of 3-s, consistent with the approach used in prior research [6, 50–52]. Then, we have generated the mel-spectrogram images from those signal segments using STFT and mel filter bins. After these segmentation and image generation steps, the resulting dataset contains 9610 and 7713 images, respectively. The repOD dataset provided 4339 images of HC and 5271 images of ScZ, while the basic sensory dataset ended up with 3102 HC and 4611 ScZ images. The dimensions of those images are 224×224 pixels and are then used as input for the CNN model. The same methodology is applied to several brain regions to produce mel-spectrogram images using the channel data from those areas, ensuring consistency and facilitating analyses of the brain lobes. The studies are performed on a PC with a configuration setup that included 128 GB of RAM and a 16 GB of graphics memory, providing sufficient resources for effective processing and analysis. We have performed 100 epochs and a training batch size of 32 to train the CNN model.

### Results

This study has proposed a comprehensive framework to identify the essential brain lobes implicated in EEG signal data for the identification of ScZ. The main aim was to assess and compare the effectiveness of this framework in differentiating between individuals with ScZ and HC. The framework underwent evaluation through a 10-fold CV process to ascertain the robustness and generalizability of the findings. Then, we applied LIME, SHAP, and Grad-CAM for the explainability of the ScZ diagnosis. Then, the same method is followed across various brain lobes to see which regions most significantly contributed to differentiating ScZ patients from HCs in classification tests. Tables 3 and 4 show the 10-fold CV average result of the experiments for various brain lobes for the two considered datasets in this study.

**Table 3.** Brain Lobes Classification Results - RepOD dataset.

| Metrics | Full brain | Central | Frontal | Occipital | Parietal | Temporal |
| --- | --- | --- | --- | --- | --- | --- |
| Sensitivity | 99.91 ± 0.13 | 91.39 ± 1.85 | 97.65 ± 0.70 | 86.30 ± 4.76 | 91.24 ± 1.21 | 96.71 ± 1.00 |
| Specificity | 99.72 ± 0.20 | 90.91 ± 1.46 | 96.27 ± 0.65 | 70.36 ± 23.64 | 90.58 ± 1.34 | 95.99 ± 0.94 |
| Precision | 99.77 ± 0.16 | 92.45 ± 1.04 | 96.95 ± 0.56 | 79.88 ± 7.90 | 92.16 ± 1.16 | 96.67 ± 0.89 |
| F1 | 99.84 ± 0.12 | 91.90 ± 0.90 | 97.29 ± 0.33 | 82.47 ± 3.49 | 91.69 ± 0.75 | 96.69 ± 0.62 |
| Accuracy | **99.82 ± 0.13** | 91.19 ± 0.80 | **97.02 ± 0.34** | 79.30 ± 7.61 | 90.94 ± 0.74 | 96.37 ± 0.66 |
| FPR | 0.28 ± 0.20 | 9.09 ± 1.46 | 3.73 ± 0.65 | 29.64 ± 23.64 | 9.42 ± 1.34 | 4.01 ± 0.94 |

Table notes: Performance metrics for brain lobe classification across different brain regions. Bold values indicate the highest accuracy scores. Results show mean ± standard deviation across multiple runs.

**Table 4.** Brain Lobes classification Results - basic sensory kaggle dataset.

| Metrics | Full brain | Central | Frontal | Occipital | Parietal | Temporal |
| --- | --- | --- | --- | --- | --- | --- |
| Sensitivity | 98.89 ± 0.31 | 82.08 ± 2.04 | 91.24 ± 2.40 | 78.49 ± 4.58 | 79.83 ± 2.58 | 81.71 ± 4.03 |
| Specificity | 97.43 ± 1.13 | 54.57 ± 4.90 | 83.19 ± 3.53 | 53.02 ± 5.46 | 51.65 ± 5.22 | 61.05 ± 5.26 |
| Precision | 98.31 ± 0.66 | 72.93 ± 2.12 | 89.07 ± 1.62 | 71.37 ± 2.09 | 71.11 ± 2.54 | 75.82 ± 1.64 |
| F1 | 98.60 ± 0.33 | 77.20 ± 1.39 | 90.11 ± 1.17 | 74.68 ± 2.40 | 75.15 ± 1.34 | 78.59 ± 1.87 |
| Accuracy | **98.31 ± 0.41** | 71.04 ± 1.77 | **88.03 ± 1.45** | 68.33 ± 1.66 | 68.46 ± 1.92 | 73.47 ± 1.57 |
| FPR | 2.57 ± 1.13 | 45.43 ± 4.90 | 16.81 ± 3.53 | 46.98 ± 5.46 | 48.35 ± 5.22 | 38.95 ± 5.26 |

Table notes: Performance metrics for brain lobe classification across different brain regions. Bold values indicate the highest accuracy scores. Results show mean ± standard deviation across multiple runs.

Table 3 highlights the performance of our proposed framework for the ScZ vs. HC classification on the repOD dataset. It reports that, when considering the full brain region, we achieved a notable accuracy of 99.82%, with a minimal standard deviation of *±* 0.13. For the Kaggle dataset in the table 4 shows, the framework performed slightly lower but still achieved a strong accuracy of 98.31%, with a standard deviation of *±* 0.41, which is one of a least explored datasets for ScZ diagnosis. We also examined the framework’s performance across individual brain lobes to determine how specific regions contributed to the overall classification accuracy. The accuracy varied among the five brain lobes, with the Occipital lobe showing the lowest performance at 79.30 *±* 7.61 and 68.33 *±* 1.66 on both datasets. Then, it was followed by the Parietal lobe at 90.94 *±* 0.74 and 68.46 *±* 1.92 on both occasions. The Central lobe achieved an accuracy of 91.19 ± 0.80 and 71.04 *±* 1.77, which is slightly better than the Parietal lobe, while the temporal lobe performed better at 96.37 *±* 0.66 and 73.47 *±* 1.57 both times. Notably, the frontal lobe yielded the highest accuracy at 97.02 *±* 0.34 and 88.03 *±* 1.45, as provided on the different occasions.

To improve our understanding of the performance of several brain lobes across different evaluation metrics, we visually contrasted sensitivity, specificity, precision, and accuracy for each lobe with error bars. Fig 5, 6, 7, and 8 illustrate these variations, offering an accurate representation of the performance across the two datasets. Since we constructed the performance metrics on two datasets, it is important to visualize the contribution of each brain lobe across the datasets. These visualizations clarify the variety in performance measures throughout the brain lobes.

**Fig 5.**
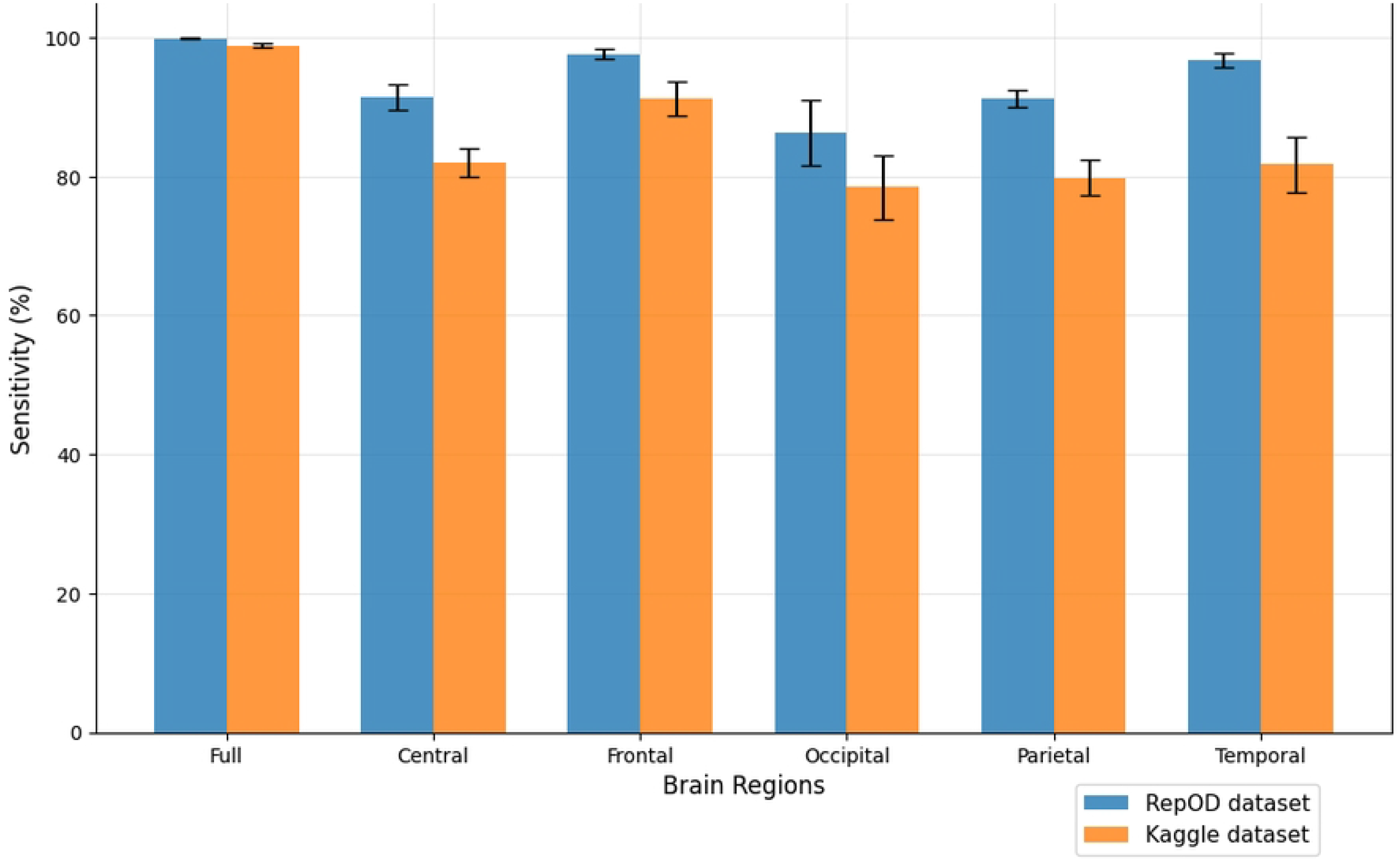
Sensitivity comparison across different brain regions and dataset.

Fig. 5 illustrates a lobe-specific comparison of sensitivity values for both datasets in ScZ vs HC classification. Sensitivity in this context denotes a brain region’s ability to accurately identify individuals with the disease, hence reducing false negatives. The entire brain region exhibits an average sensitivity of 99.91% in the repOD dataset, demonstrating the brain’s effectiveness in detecting ScZ Disease. In contrast, for the Kaggle dataset classification, it exhibits an average sensitivity of 98.89%, suggesting it is a commendable performance at recognizing patients with ScZ disease. The frontal lobe exhibits the highest sensitivity among the five brain lobes, with a value of 97.65% and 91.24% both times. This sensitivity is almost as close to that of the entire brain region, emphasizing the frontal lobe’s specific efficacy in recognizing patients with ScZ disease. The parietal and central lobes similarly show considerable sensitivity, with 91.24% and 91.39%, respectively, providing them with moderate possibilities for disease classification. On the contrary, it remains a similar pattern in demonstrating comparatively lower sensitivity scores with 79.83% and 82.08% on the Kaggle dataset. The occipital lobe, conveys a poor performance with sensitivity of 86.30% and 78.49% in both times. The temporal lobe has the second-best performance, achieving an average sensitivity of 96.71% in the repOD dataset, while there is a big fall to 81.71% in the Kaggle dataset. It exhibits impressive results but is inferior to the frontal lobe in terms of sensitivity. The frontal lobe is prominent in ScZ disease detection, where the temporal lobe produced inconsistent performance.

Fig. 6 provides a detailed comparison of the specificity percentages across different brain regions. Specificity, in this context, refers to the ability of a brain region to correctly identify those without the disease, or in other words, its ability to avoid false positives. In the repOD dataset, the frontal region stands out with the highest specificity of 96.27%, which suggests it is particularly effective at identifying HCs and distinguishing them from individuals with ScZ. The full region shows high specificity at 99.72%, confirming its broad capability to differentiate between ScZ and HC. However, the occipital region does not perform well in this task, with a notably low specificity of just 70.36% and 53.02% both occasions. This indicates that the occipital region is not very effective at ruling out ScZ in individuals who do not have the disease, leading to a high rate of false positives. Both central and parietal lobe show moderate performance levels with the specificity of 90.91% and 90.58% which are almost close, but not as high a value as frontal and temporal. Again, like sensitivity, the temporal region showed the second-highest specificity score of 95.99%, which illustrates its particular significance in the prediction. But, in the Kaggle dataset, the specificity is a bit lower with 81.71%, which suggests the temporal region has inconsistent specificity. Some regions, such as the frontal and full regions, are consistently reliable in distinguishing ScZ vs HC. In contrast, others, like the occipital region, may require further evaluation or different strategies depending on the context.

**Fig 6.**
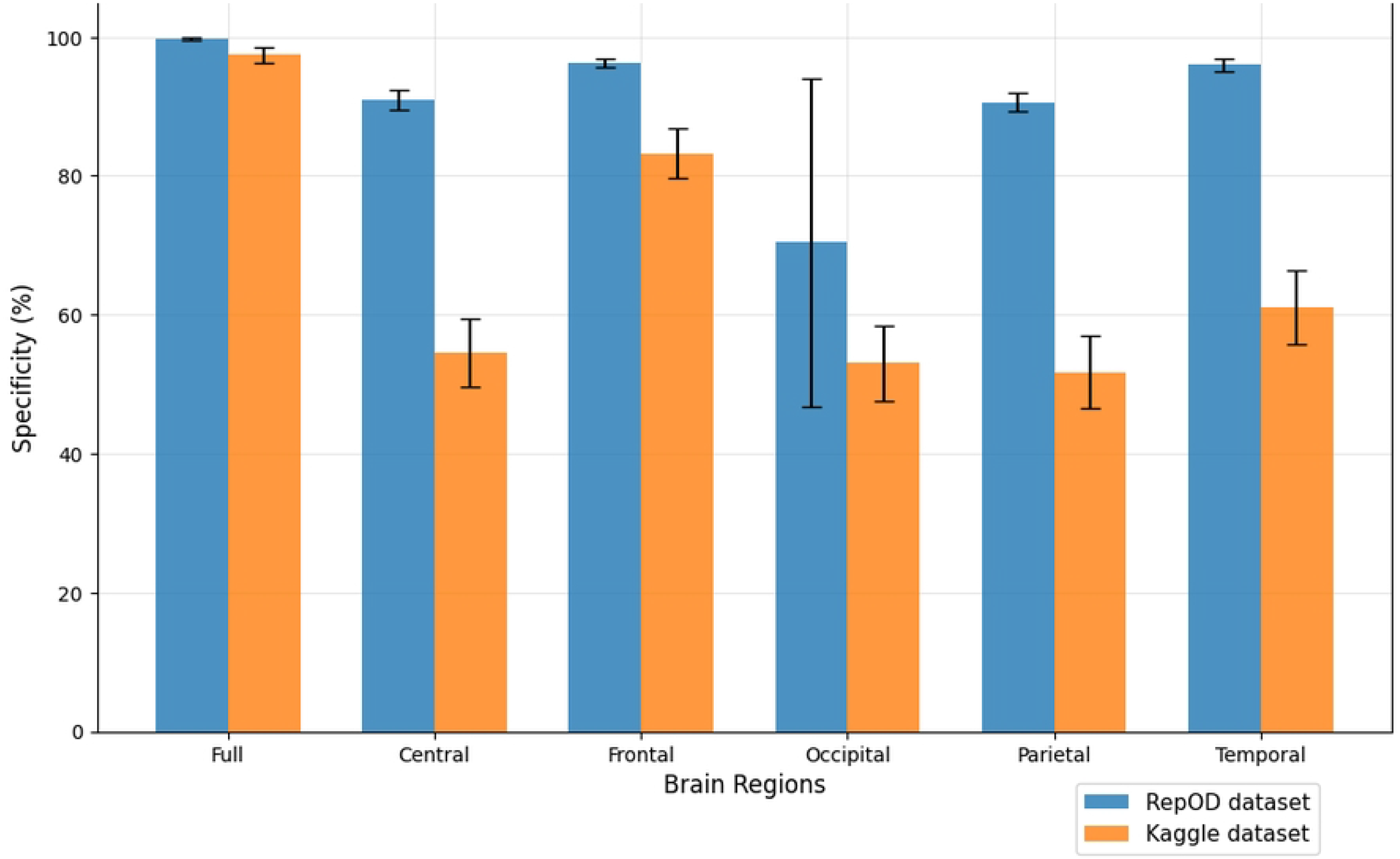
Specificity comparison across different brain regions and dataset.

Fig. 7 provides a comprehensive illustration of the precision percentages across various brain regions. In the repOD dataset, the full region and frontal region both show high precision, achieving 99.77% and 96.95%, respectively. Temporal showed almost close values compared to these with 96.67%, but it reduces significantly in the Kaggle dataset. However, the occipital region demonstrates significantly lower precision at 79.88%. The central and parietal regions show moderate precision values of 92.45% and 92.16%, like the other metrics in this study, they again show almost similar percentages. In the Kaggle dataset, it is observed that the precision of all the brain regions follows the hierarchy of the repOD dataset. The frontal region again leads with the highest precision, and the occipital region continues to show the lowest precision. Figure 7 demonstrates that the variation in precision across the various lobes between the two groups is consistent.

**Fig 7.**
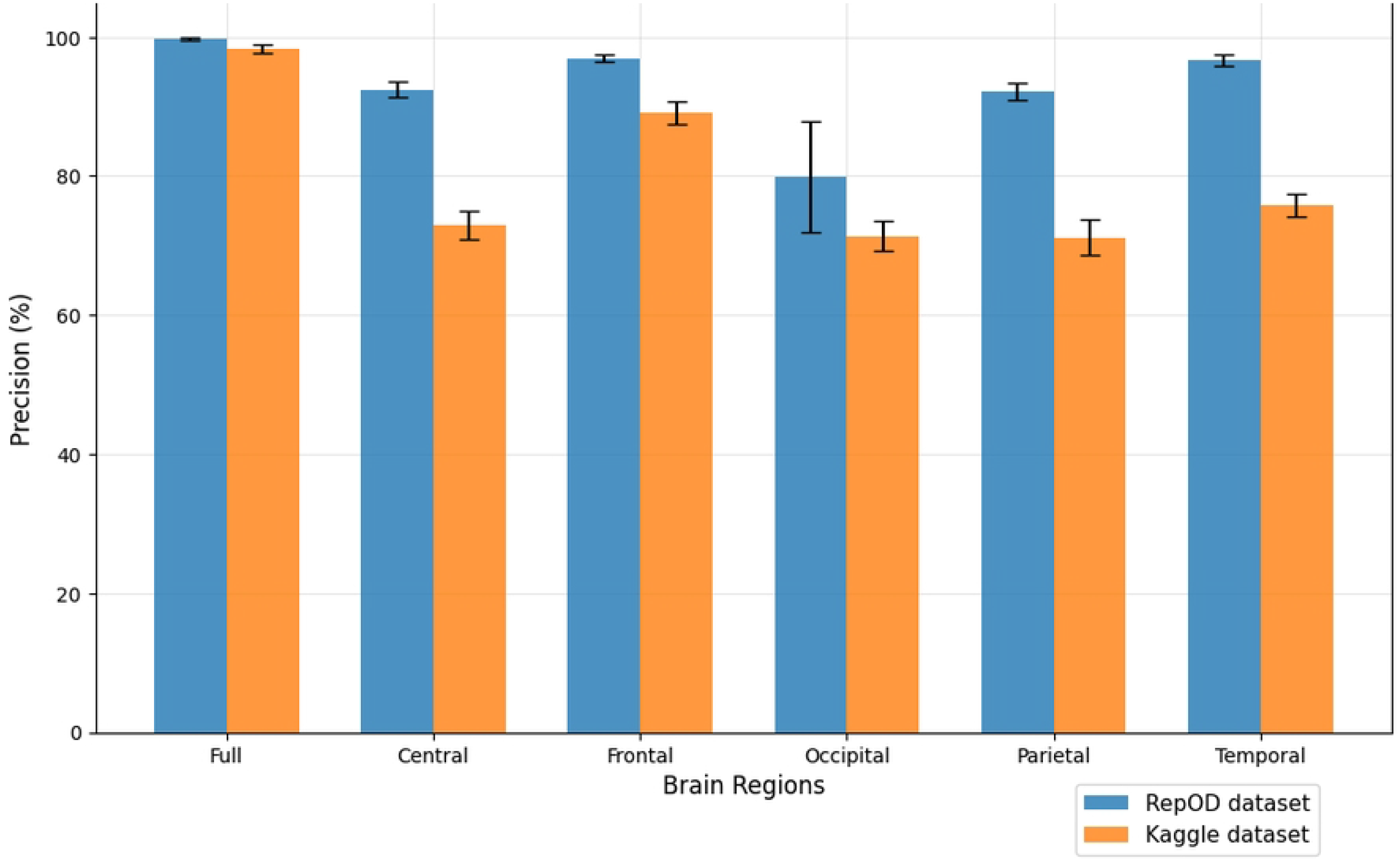
Precision comparison across different brain regions and dataset.

Fig. 8 provides a comprehensive comparison of accuracy across different brain regions in classifying ScZ vs. HC. It is demonstrated that the whole brain region achieves an accuracy of 99.82% and 98.31% in both datasets. Among the other regions, the frontal lobe demonstrates strong performance, achieving an accuracy of 97.02% and 88.03% on both occasions. In contrast, the Occipital brain region shows the lowest performance, with an accuracy of just 79.30% and 68.33% in both times. The temporal lobe shows the second-highest accuracy among the lobes, with 96.37% accuracy, but it is notably reduced in the Kaggle dataset to 73.47%. Then, the central and parietal lobes yield more moderate accuracy levels with 91.19% and 90.94% respectively, showing that these regions play a reasonable critical role in distinguishing ScZ vs. HC compared to others. In the Kaggle dataset, these two show an accuracy of 71.04% and 68.46%. This analysis underscores the variability in diagnostic accuracy across different brain regions, highlighting the superior performance of the frontal regions in both classification tasks. Besides, the temporal lobe showed a better accuracy in repOD dataset, aiding significant values but not stable ones. It is clearly seen that the results orientation among the lobes is almost similar. Furthermore, the consistency of these findings across both datasets and conditions reinforces the significance of specific brain regions, particularly the frontal lobe.

**Fig 8.**
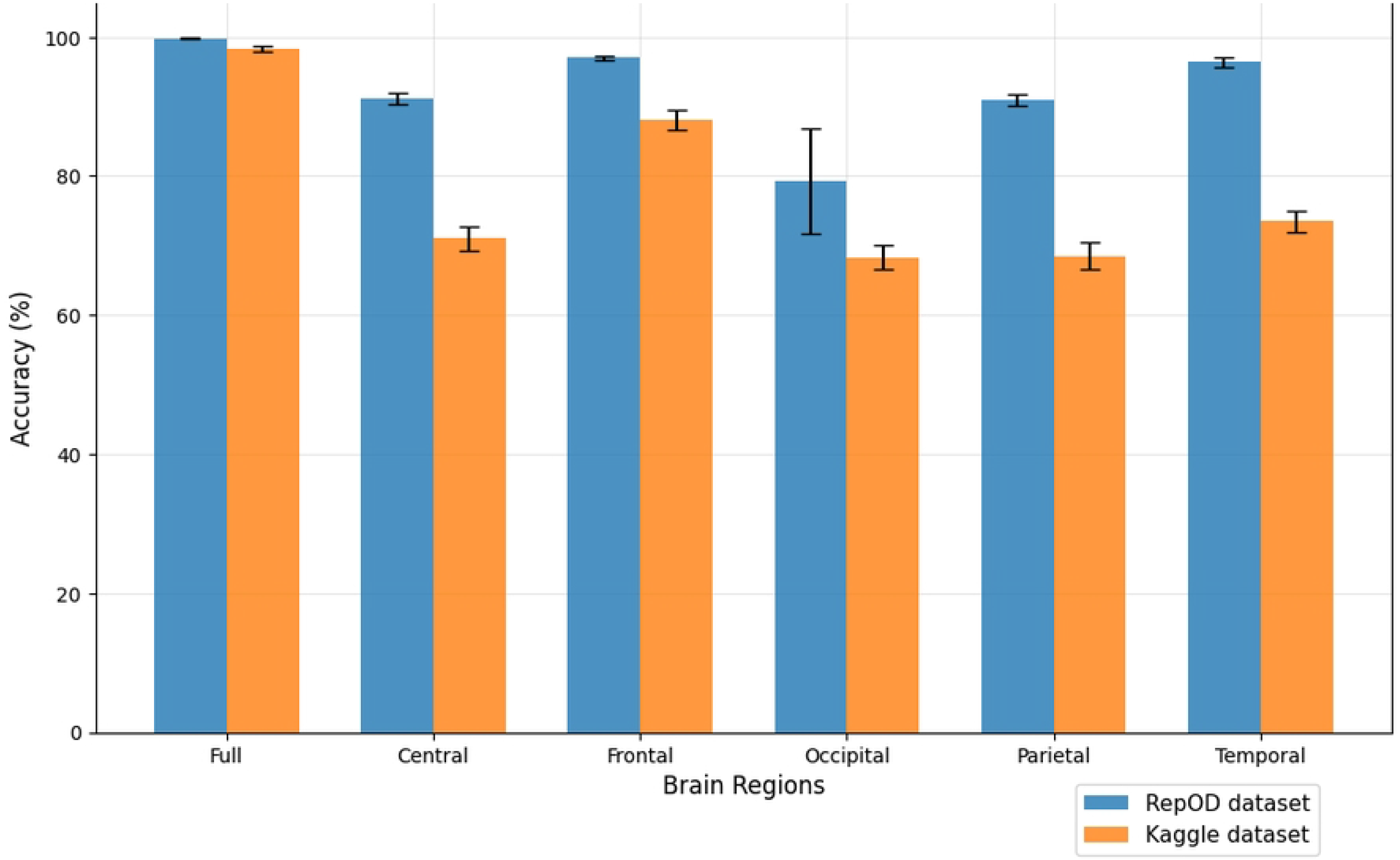
Accuracy comparison across different brain regions and dataset.

The ROC curve is an illustration that plots the true positive rate on the y-axis and the false positive rate on the x-axis. The curve for the full brain region, represented by the purple line in Fig. 9, demonstrates a nearly flawless classification performance. The strong elevation and position in the top-left corner of the graph signify a classifier that attains both high sensitivity (accurately identifying ScZ) and a minimal false positive rate (reducing erroneous identification). The frontal region, shown by the blue line, is close to the curve for the full brain lobe. This indicates that the frontal lobe data are highly informative for distinguishing. Other brain regions, including the parietal (cyan) and central (orange), showed moderate performance. Despite maintaining relatively high true positive rates, the performance variability indicates that these regions may not be as dependable for differentiating ScZ from HC as the entire brain or frontal region. Additionally, the temporal (olive) regions show better performance compared to the other regions, as they are closer to the frontal curve. Conversely, the occipital region (shown by the red line) demonstrates significantly inferior performance. This line diverges from the top-left corner, signifying a higher false positive rate. Overall, the ROC curve in Fig. 9 indicates that the entire brain and frontal regions provide optimal performance. The temporal region demonstrates significant significance; although other regions show potential, their performance varies, with the occipital region notably under performing in accurate classifications.

**Fig 9.**
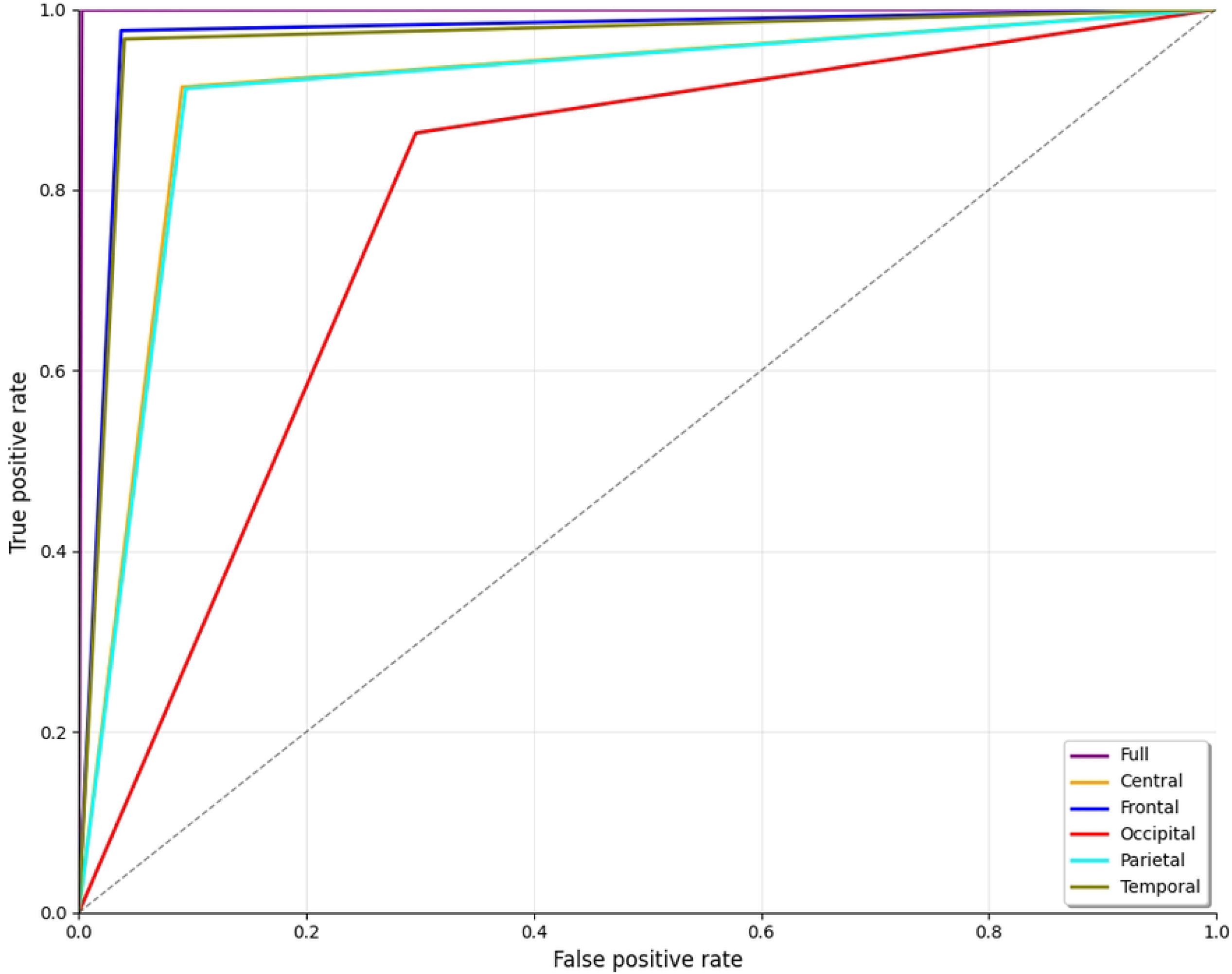
ROC curve for ScZ vs. HC classification for RepOD dataset.

The ROC curve shown in Fig. 10 illustrates the classifier’s performance across different brain regions for distinguishing ScZ vs HC for the Kaggle dataset. Here again, the frontal region represented by the blue line also shows excellent performance, with a curve closely following the full brain analysis. Other brain regions, including the central region (orange) and parietal(cyan) regions, demonstrate balance performance. In contrast, the occipital region (shown by the red line) has poor outcomes in comparison. The deviation from the top-left corner of the graph indicates a high false positive rate and reduced sensitivity. A noticeable thing is that the temporal (olive) regions showed better performance compared to the other regions in the repOD dataset; this time, it shows a significantly lower performance, like the central and parietal lobes. This line position indicates that the temporal lobe is significant to overall outcomes, with areas providing vital information, although less accurately than the entire brain or frontal regions.

**Fig 10.**
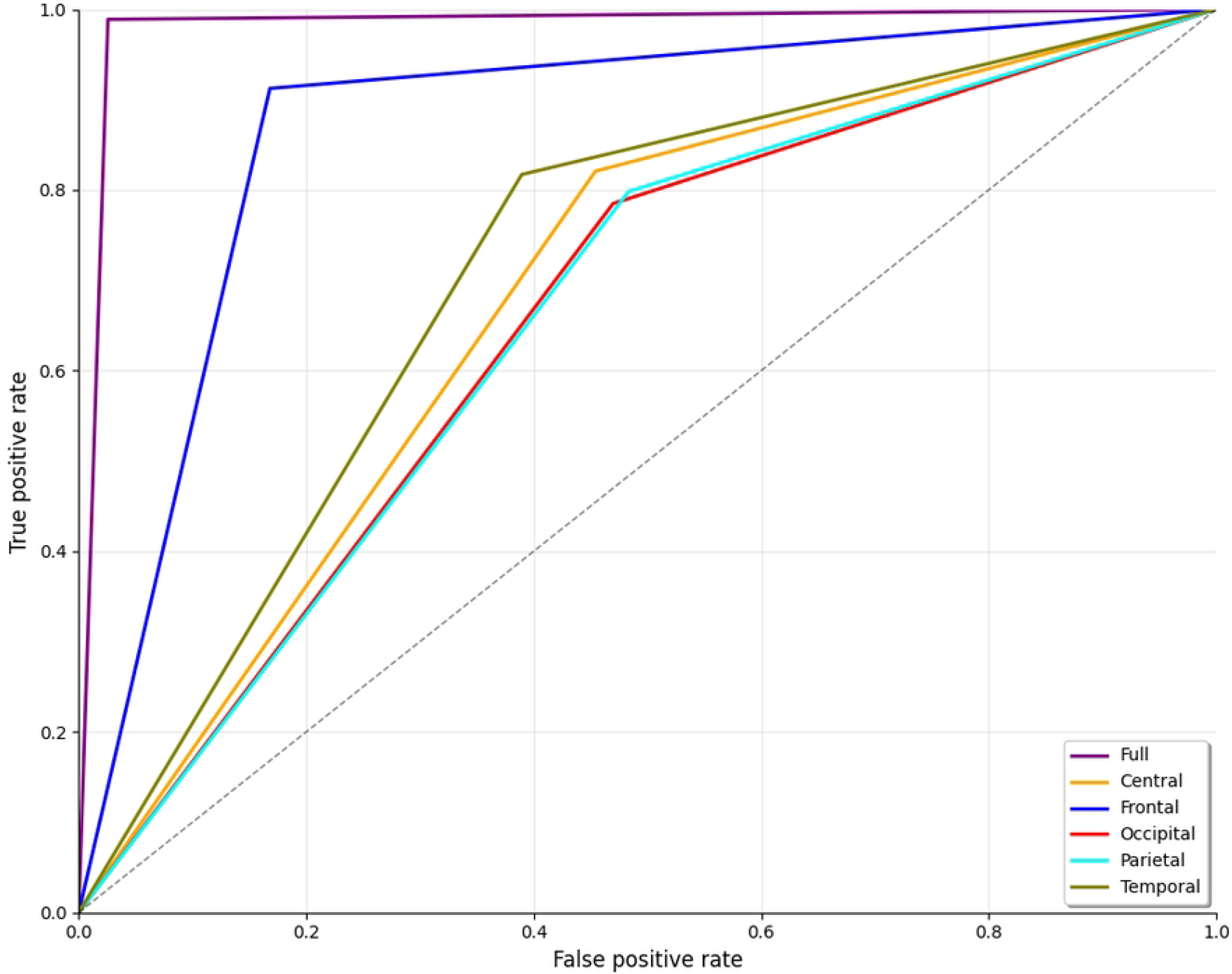
ROC curve for ScZ vs. HC classification for kaggle basic sensory task dataset.

### Explainability analysis

In recent years, Explainable AI (XAI) has evolved as a tremendous method for model explainability. XAI has become essential in understanding, validating, and trusting machine learning models, especially in sensitive domains like healthcare. In our study we have introduced three popular XAI techniques to explain the result outcomes.

### LIME

LIME (Local Interpretable Model-agnostic Explanations) basically works by approximating the complex CNN’s behavior locally with an interpretable, simpler model, such as a linear regression, around a specific prediction instance. LIME is applied to individual mel-spectrogram images to generate heatmap-like explanations that reveal which frequency bands and time windows most influence the model’s predictions. This detailed analysis often shows that certain frequency rhythms, like altered gamma or theta bands, consistently stand out as important markers for identifying ScZ. Essentially, LIME breaks the mel-spectrogram into segments and visually indicates which parts of the image contribute positively or negatively to the model’s decision, making the prediction more understandable and transparent.

We created LIME visualizations for the full brain region as well as for five individual brain regions. For two different datasets, we have done the LIME visualizations. The LIME explanations for all five brain regions, along with the full region, are shown in Fig. 11 and 12 for the repOD dataset. Fig. 11 show healthy images for the first image after segmentation, and Fig. 12 shows the corresponding first image of the ScZ patient image, which was classified by our CNN model across different brain regions. It is clearly observed which portion of the image is contributing positively and negatively in the disease classification. It basically identified a region as P1, P2, P3, etc, which means they are positively contributing to the model outcomes, and N1, N2, N3, etc, that negatively contribute to the model. Here, P1 and N1 are denoting the most contributing, then P2, N2, and so on. It is seen that the positive contributing and negative contributing segments in an image are different. When one segment of a ScZ image is positively predicted as ScZ, a different portion of the image is predicted as HC. Similarly, Figures 13 and 14 demonstrate the LIME explanations for all five regions, along with the full region for the Kaggle dataset. A similar pattern is consistently seen across all LIME explanations in both datasets.

**Fig 11.**
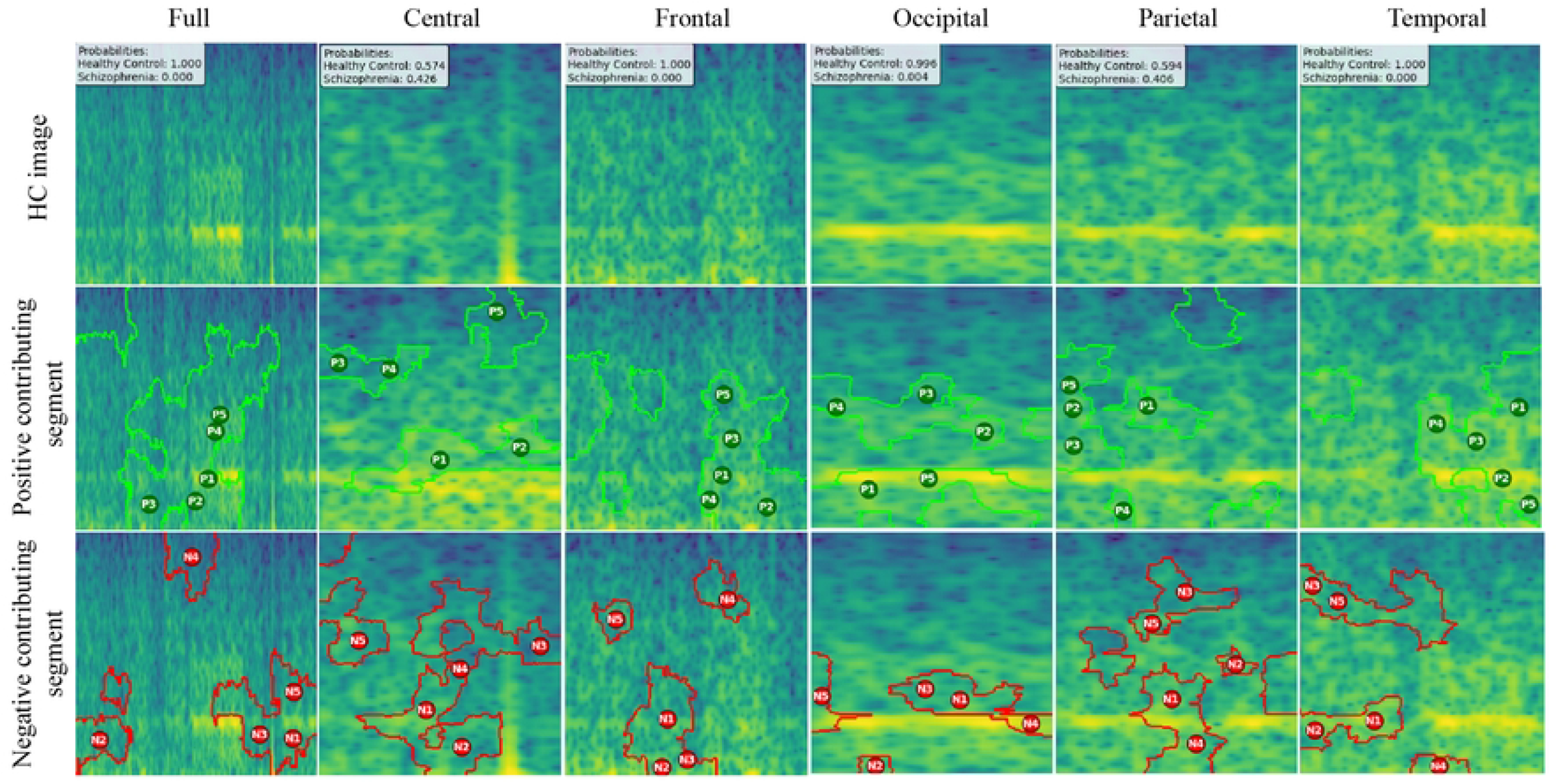
RepOD dataset LIME contributing segment in ScZ vs. HC classification for HC images in first segment for different brain regions.

**Fig 12.**
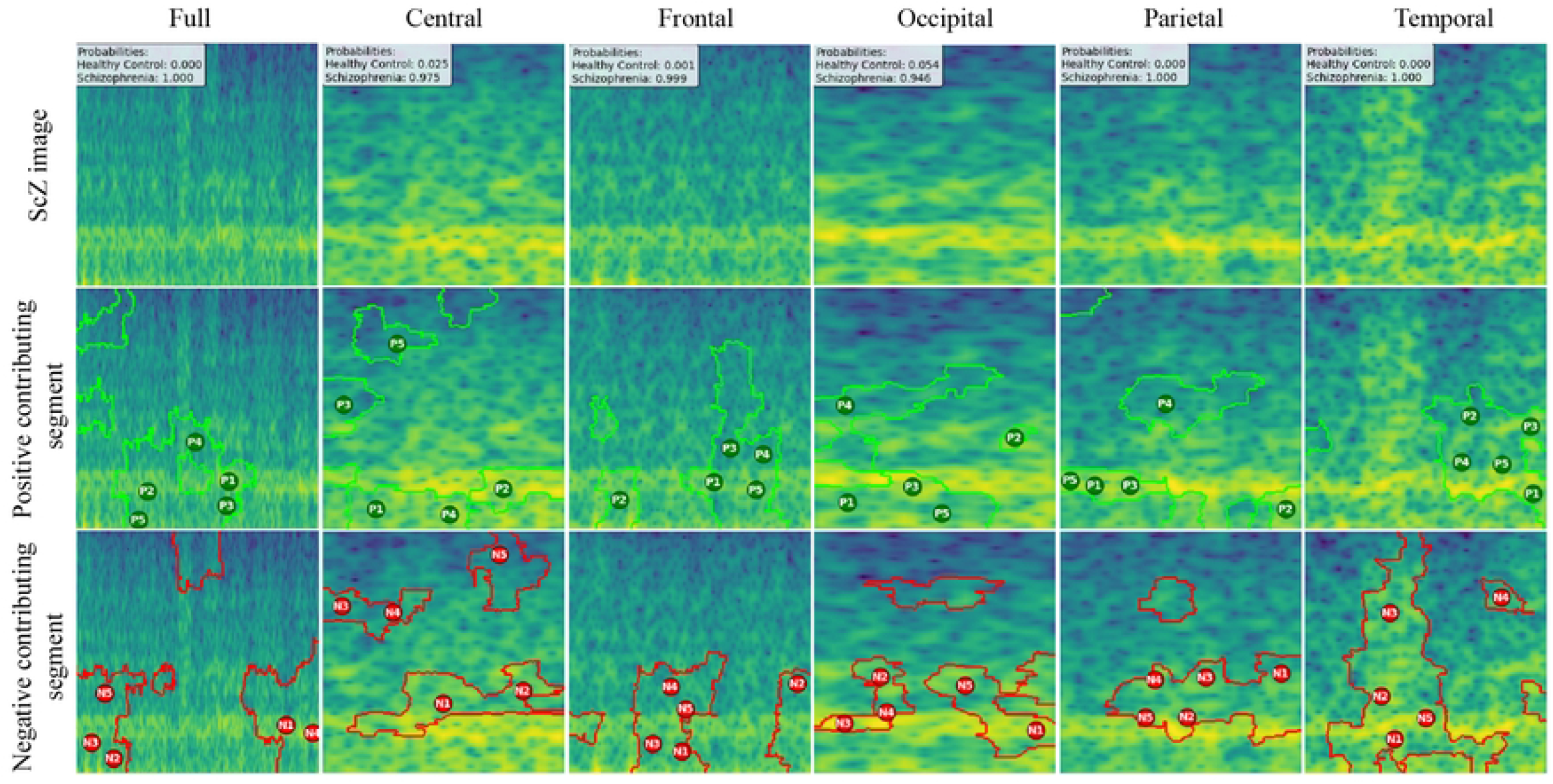
RepOD dataset LIME contributing segment in ScZ vs. HC classification for ScZ images in first segment for different brain regions.

**Fig 13.**
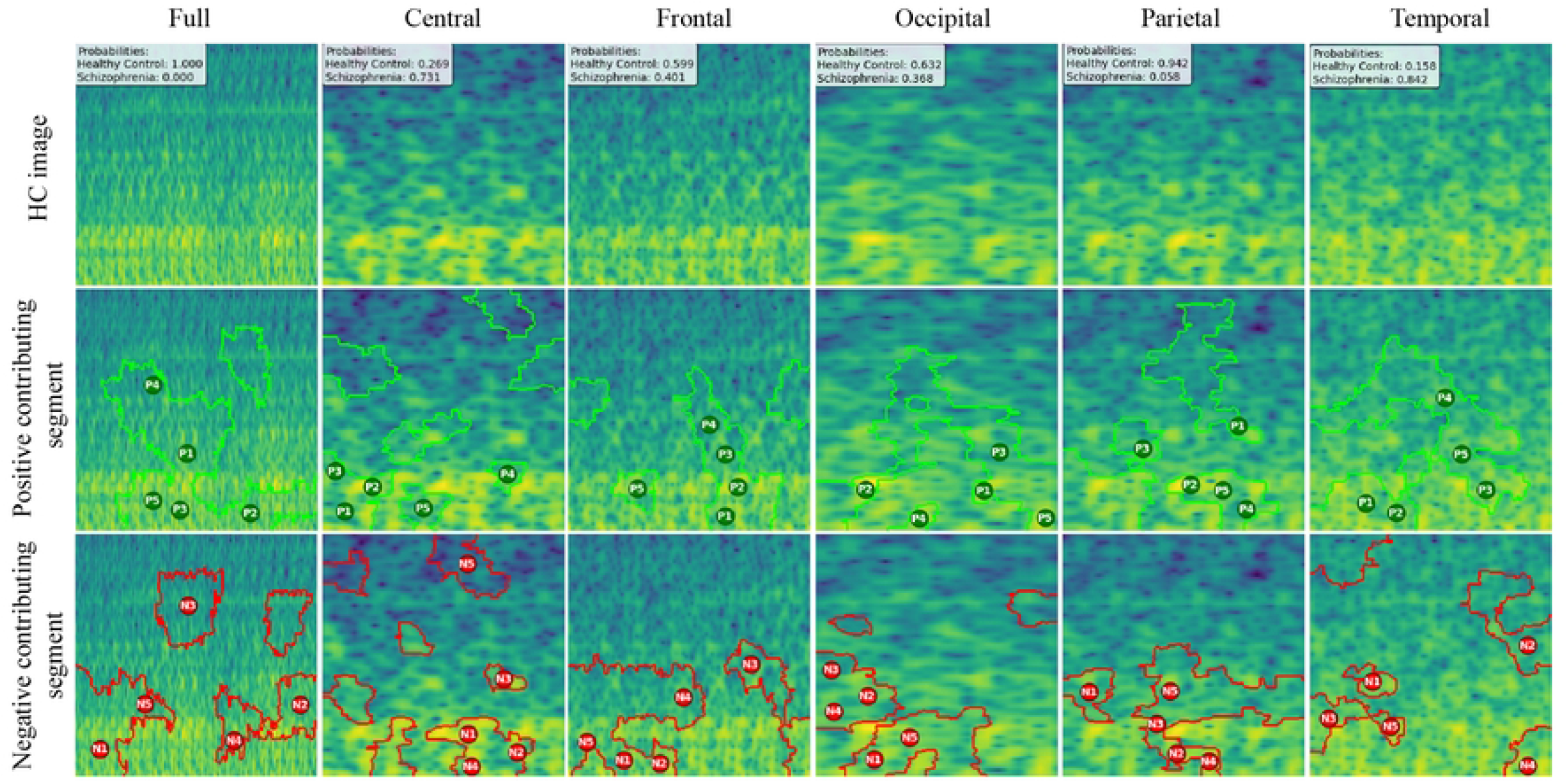
Kaggle dataset LIME contributing segment in ScZ vs. HC classification for HC images in first segment for different brain regions.

**Fig 14.**
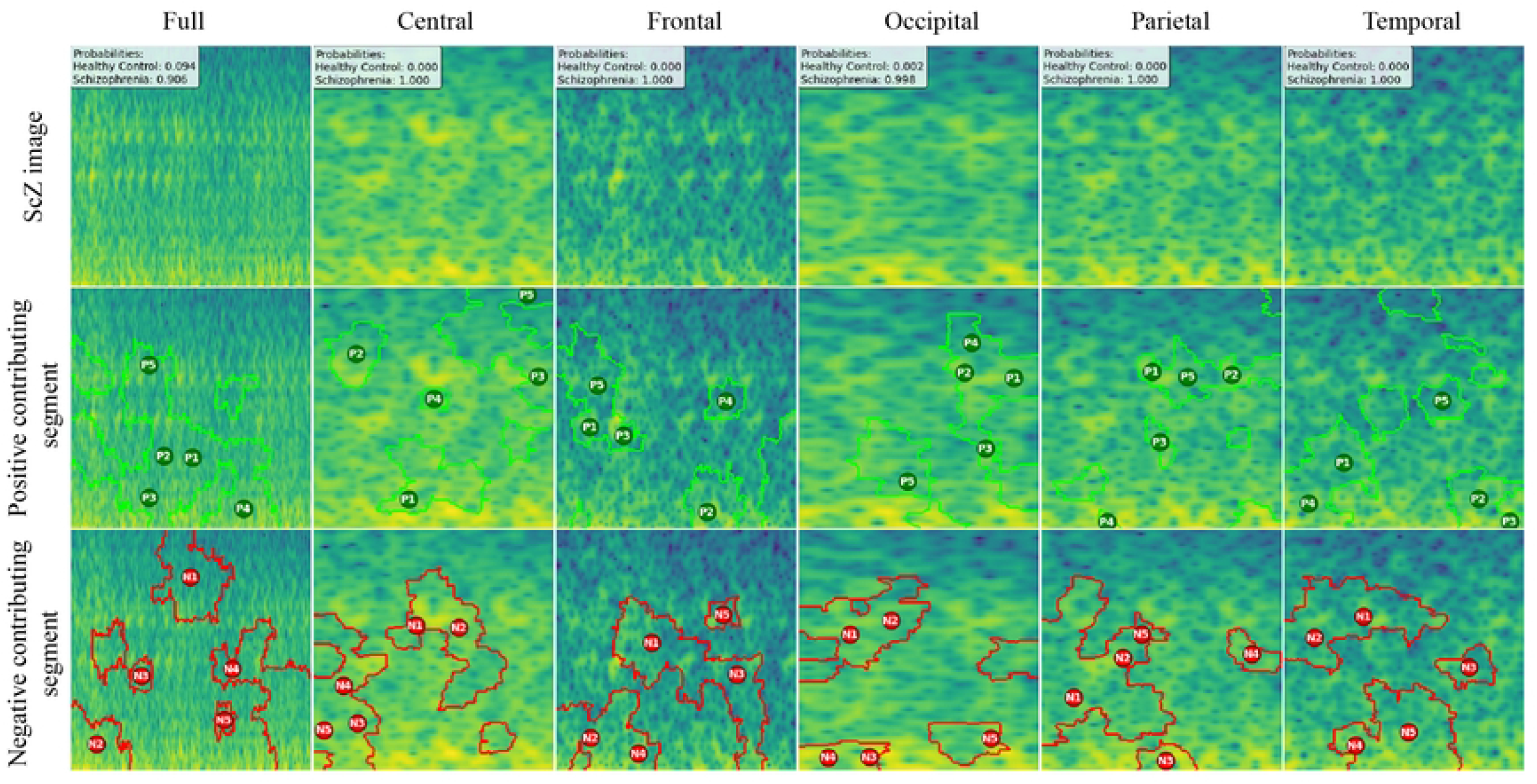
Kaggle dataset LIME contributing segment in ScZ vs. HC classification for ScZ images in first segment for different brain regions.

These findings can be understood from various perspectives. It is seen that, in the full region for repOD, the positive contributing segment in the healthy image is ranked as *P*_1_ *> P*_2_ *> P*_3_, etc, and in the Kaggle dataset is *P*_1_ *> P*_2_ *> P*_3_ etc. The contributing segment portion is almost the same in both datasets. So, it is clearly observed which portion is reasonable for diagnosis ScZ or not. This suggests that our model accurately predicts outcomes and identifies important features effectively. Since the input images are time-frequency mel spectrograms with time on the x-axis and frequency on the xis (filtered between 0.5 and 45 Hz), we notice that segments contributing to the ScZ classification are mainly located in the lower frequency bands in both datasets. These correspond to the delta and theta bands, which are dominant during deep, dreamless sleep and are also present in infants. Theta bands are associated with deep relaxation and meditation. This insight may help clinicians in diagnosing and treating ScZ. On the other hand, the HC images were mainly associated with alpha and beta bands, as seen in the image prediction, contributing positively to a higher portion of the image. These observations provide helpful information for doctors in their diagnostic decision-making process.

From the earlier results, it was evident that the frontal brain region provided the highest accuracy in classifying ScZ, while the occipital lobe showed the lowest accuracy.

A clear pattern emerges from these images: the segments that contribute positively and negatively to the model’s predictions in the frontal region are very similar and closely align with those observed in the full brain region’s LIME explanations. This close alignment suggests that the frontal region carries much of the predictive information that drives the model’s success, supporting its importance in ScZ diagnosis. The high accuracy achieved with the frontal region demonstrates that it captures critical features related to the disorder. In contrast, the key contributing segments of the occipital region’s mel-spectrogram are spatially far away from those identified in the full brain region. This finding suggests that the occipital lobe has a weaker connection to the features important for diagnosing ScZ according to our model. The frontal lobe plays a critical role in capturing neural patterns linked to ScZ, making it an influential region for model predictions. On the other hand, the occipital lobe’s contribution is minimal or even contrasting, indicating it is less relevant to the diagnosis. From a clinical perspective, practitioners can focus more on frontal lobe-specific treatment procedures when interpreting EEG data for ScZ diagnosis, potentially improving the accuracy and reliability of their assessments. At the same time, recognizing that the occipital region contributes less can help streamline diagnostic processes and avoid unnecessary emphasis on less informative brain areas. Ultimately, these insights can guide targeted treatment strategies and support better decision-making in managing ScZ.

### SHAP

SHAP (SHapley Additive exPlanations) is a technique used to explain the predictions of machine learning models by assigning a contribution value to each feature. SHAP basically ranked the feature importance of the model which is why its transparency and fairness have made SHAP a prevalent tool for decision-making in fields like healthcare.

We applied SHAP only for the full region to identify the most important features in the model’s decisions. As we filtered the image from 0.5 to 45 Hz. Fig. 15 and 16 shows important features from the full brain region, in classification tasks, ScZ vs HC. It produces feature importance by ranking time segments and frequency range. It is observed that the full brain SHAP ranking has similar feature importance in both datasets.

**Fig 15.**
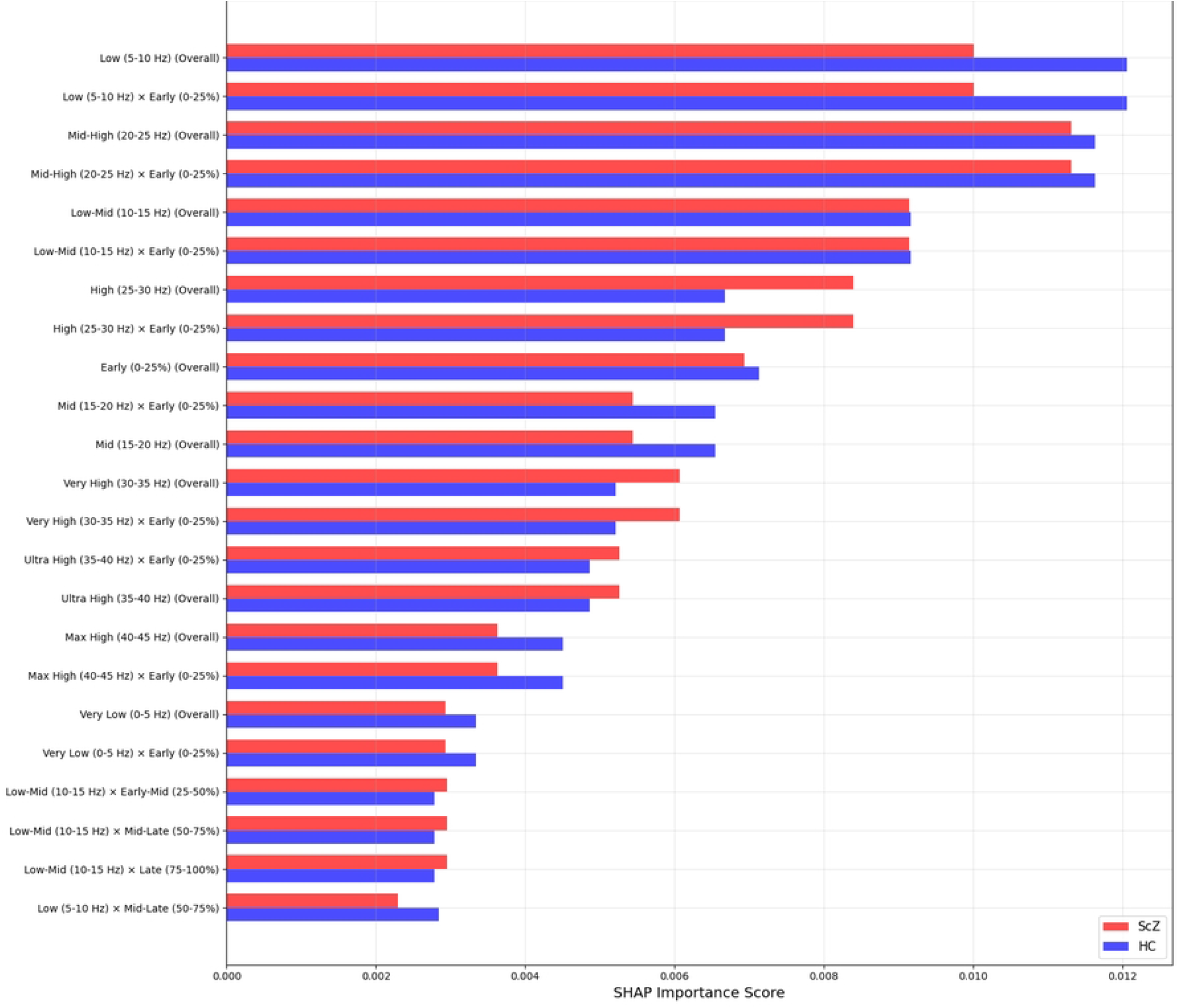
SHAP feature importance rankings for repOD dataset for the full region.

**Fig 16.**
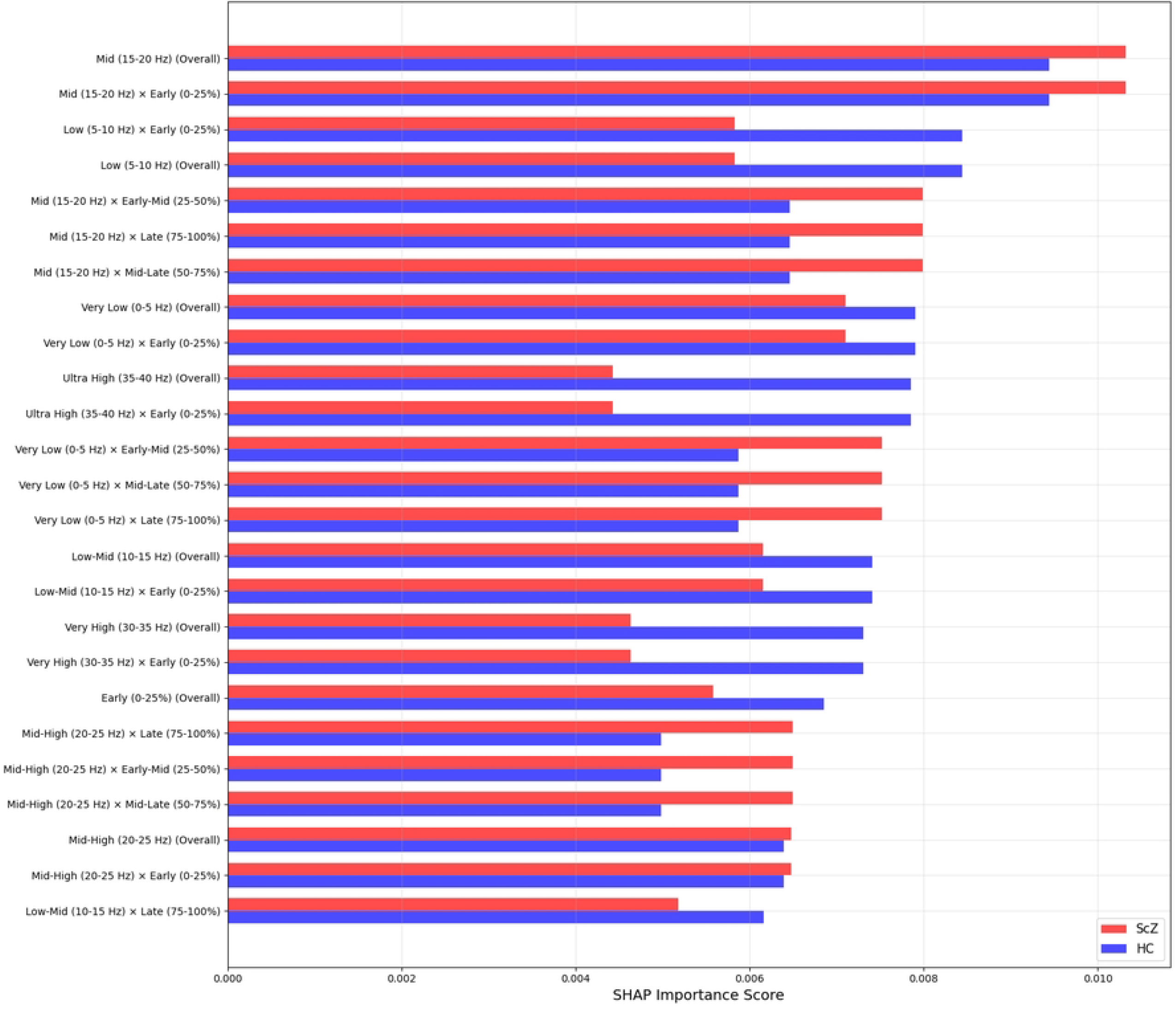
SHAP feature importance rankings for Kaggle basic sensory task dataset for the full region.

It is clearly visible in the Fig 15 that in the repOD dataset, the full region with the most influential frequency band spans the mid-level (20-25 Hz) and low-level (0-5 Hz) frequency ranges for ScZ diagnosis. Conversely, for HCs, the range is also the same. LIME visualizations also confirm this range. Then, a slightly varied range is identified in the basic Kaggle dataset in Fig. 16. It is visible that, for ScZ, the range is between 15-20 Hz and 0-5 Hz. Furthermore, for HCs, the range lies mostly between 15-20 Hz and 5-10 Hz. These are nearly similar in both datasets and also establish the outcomes of other XAI techniques

### Grad-CAM

Grad-CAM (Gradient-weighted Class Activation Mapping) is a widespread approach utilized to visualize and analyze the decision-making process of CNNs by spotlighting which regions of an input image were most influential for a particular prediction.

To identify the most influential mel-spectrogram features in the model’s decisions, we applied the Grad-CAM method only for the full region. Fig. 17 presents a sample HC and ScZ mel-spectrogram images from the full brain lobe, along with its corresponding Grad-CAM visualizations. It is observed that the full brain Grad-CAM image overlay has significant regions with a high heatmap area on both occasions.

**Fig 17.**
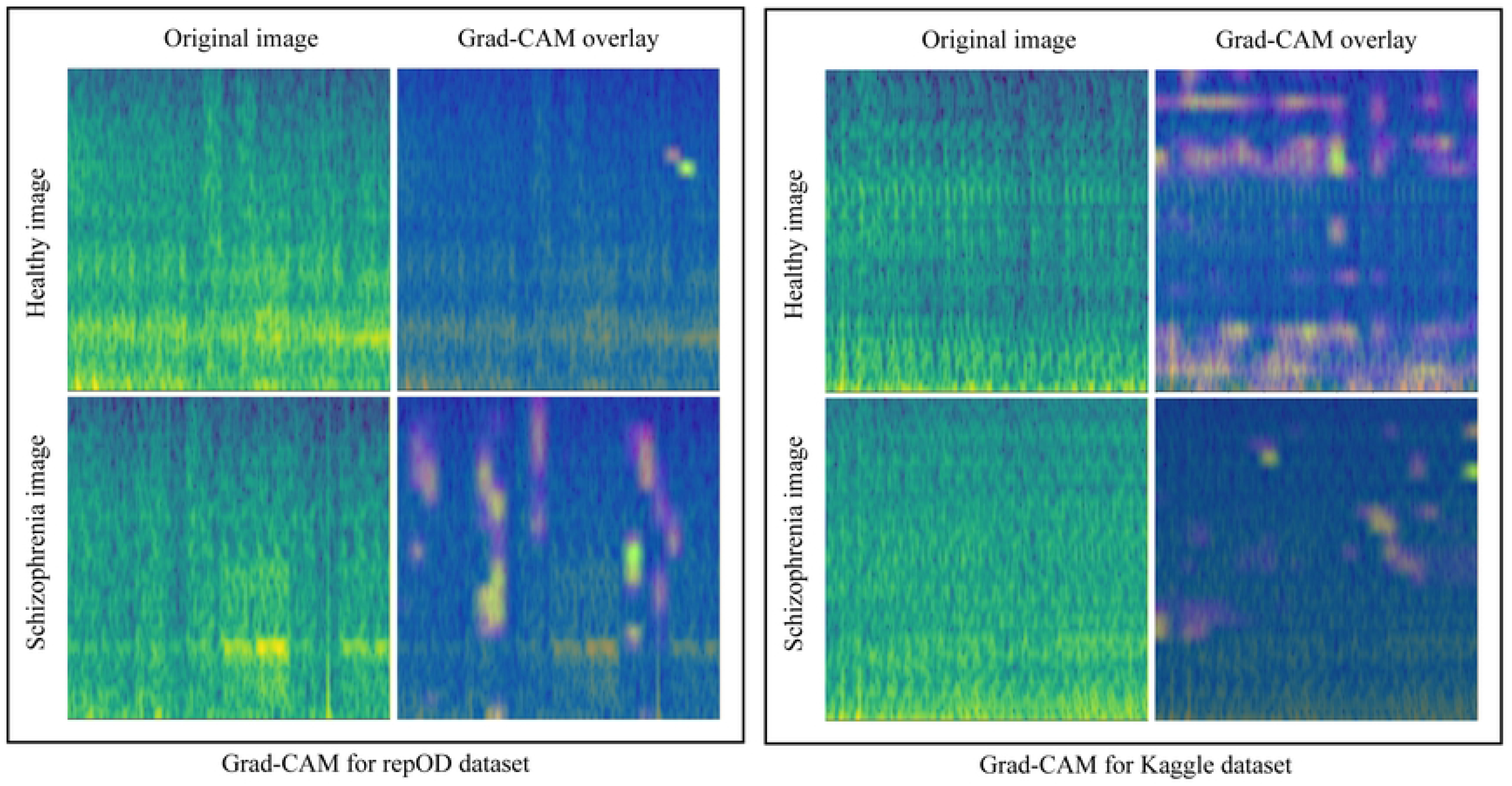
Grad-CAM for the repOD dataset and Kaggle basic sensory task dataset for the full region.

## Discussion

This study presents an approach that combines an STFT-based mel-spectrogram image and a CNN to identify critical brain lobes as biomarkers for distinguishing ScZ from HC using EEG data. At first, a butterworth band-pass filter of 0.5-45 Hz is used to clean up the EEG signals by eliminating the presence of noise and other distortions. The signals were then segmented into three-second (3s) time frames, according to the established study [6, 38, 45, 51]. The EEG channels were grouped according to their corresponding brain lobes to facilitate targeted analysis. The frontal, central, temporal, parietal, and occipital lobes are all part of these groupings. Two separate EEG datasets were used to assess the proposed architecture. For the kaggle dataset, which had 64 channels, 19 channels were chosen depending on the location they were in the International 10-20 electrode position system. Using STFT, the EEG signals were then turned into mel spectrograms, which showed brain activity in terms of time and frequency mel bins. We then put these mel-spectrograms into a CNN model to classify them. We classified the mel spectrograms from each brain lobe and the entire EEG channel set independently to see how well each one worked for finding ScZ. The findings, as shown in Table 3 and Table 4, suggested that the method worked well for accurately classifying ScZ from HC. It got a classification accuracy of 99.82% with the full brain area and 98.31% with the Kaggle dataset. The frontal lobe demonstrated the highest accuracy in both datasets; conversely, the occipital lobe has the lowest, consistent with the discovery of lobe-specific biomarkers for the detection of ScZ.

Each brain region uniquely contributes to the diagnostic process, and acknowledging these contributions enhances the accuracy and reliability of EEG-based ScZ detection [54, 56–58].

- The frontal lobe is critical for executive functions, which include higher-order cognitive processes such as decision-making, problem-solving, planning, and regulating behavior. In patients with ScZ, EEG recordings from the frontal region often show altered brain activity, specifically an increase in beta power and a decrease in theta and delta power. These patterns are believed to reflect impairments in cognitive control and executive functions, which are central to the symptoms of ScZ. The changes in these brain wave frequencies suggest a disruption in the mental processes required for effective functioning. Our study aligns with these findings, as we achieved the highest prediction accuracy when using EEG data from the frontal lobe across both datasets. These insights highlight the importance of targeting the frontal lobe in both diagnosis and therapeutic approaches for ScZ.
- The temporal lobe, especially the hippocampus, is a bit critical for memory and cognitive skills, both of which are profoundly impacted in ScZ. EEG studies have indicated that ScZ patients often suffer lower alpha and beta power in the temporal areas, combined with increased theta and delta activity. Such changes in brain wave patterns reflect neurological impairment and are critical for early identification, as they are strongly linked to the basic symptoms of ScZ. We noticed that using EEG data from the temporal region offer the second-best classification accuracies for these two datasets, showing the one major significance of temporal lobe activity in distinguishing between ScZ and HC.
- The parietal lobe is essential for the integration of sensory information and for the facilitation of spatial navigation, taste, texture, and temperature. In patients with ScZ, EEG data from the parietal region frequently exhibit an increase in alpha activity and a decrease in theta activity. Changes in brain wave patterns indicate moderate disturbances in cognitive and spatial processing capabilities, which are not frequently compromised in ScZ disease. Our study attained a moderate classification accuracy across parietal regions investigated for ScZ classification, which is almost similar to central lobe accuracy.
- Central lobe Primary ScZ symptoms are less frequently linked to EEG signals from the sensorimotor cortex and other central brain areas. These areas may continue to show altered rhythms and other brain rhythm abnormalities, though, which might prove an indication of more widespread neural network changes in ScZs. Central EEG signals can nevertheless help provide an extensive overview of the progression of ScZ, even though they might not be as diagnostic as those from other areas. This is demonstrated in our research as well, as we used this area of the brain to get the third-highest classification accuracy.
- The occipital lobe, which is mostly responsible for visual processing, shows less prominent changes in the early stages of ScZ. However, as the condition advances, EEG investigations have revealed that patients with ScZ may have a drop in alpha activity and an increase in slow-wave activity in the occipital areas. It suggest that the occipital lobe plays a less significant role in the development of ScZ. Our findings support this hypothesis, since we noticed that there were fewer improvements in classification accuracy when occipital EEG data was used, suggesting that less consideration is given to occipital activity for early and precise ScZ diagnosis.

### Ablation experiments

To make an informed decision, we have explored various investigations into different strategies that affect the performance of models. In Table 5, it is clearly seen that the proposed STFT-based mel spectrogram achieves the highest accuracy of 99.82% with the CNN model. Subsequently, a traditional spectrogram-based CNN model yielded a closer result; however, this approach has already been explored by several researchers. On the other hand, we underwent more complex transformations, such as the Hilbert-Huang transform and continuous wavelet scalograms, which resulted in a decrease in accuracy. This suggests that the choice of image representation has a significant influence on model accuracy, and the proposed mel spectrogram approach offers a substantial improvement over conventional techniques. These findings and analyses indicate that adopting a deep learning approach with a mel-spectrogram, followed by complexity feature extraction using a CNN model, can significantly enhance the model’s performance.

**Table 5.**
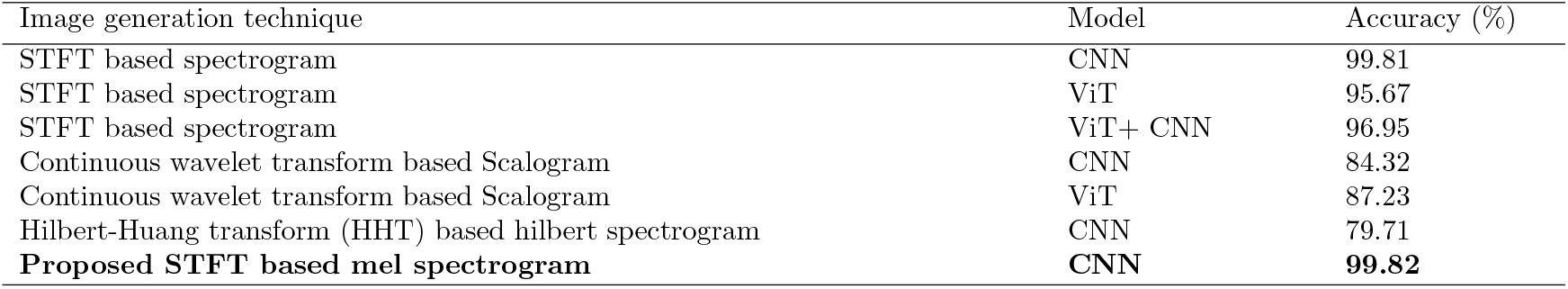
Comparison of using different image generation techniques in different models.

| Image generation technique | Model | Accuracy (%) |
| --- | --- | --- |
| STFT based spectrogram | CNN | 99.81 |
| STFT based spectrogram | ViT | 95.67 |
| STFT based spectrogram | ViT+ CNN | 96.95 |
| Continuous wavelet transform based Scalogram | CNN | 84.32 |
| Continuous wavelet transform based Scalogram | ViT | 87.23 |
| Hilbert-Huang transform (HHT) based hilbert spectrogram | CNN | 79.71 |
| **Proposed STFT based mel spectrogram** | **CNN** | **99.82** |

### Comapring with Existing Method

To provide a thorough and insightful comparison between our proposed framework and the leading state-of-the-art (SoA) methodologies, we have used the RepOD and Kaggle sensory datasets as standards. In tables 6 and 7, provide side-by-side comparison has been carefully designed to not only showcase the strengths of our framework but also emphasize the specific advancements and improvements it offers over conventional methods. This analysis serves as a critical tool for validating the effectiveness of our method and positioning it as a meaningful refinement within the current landscape of research.

**Table 6.** Comparison with existing studies on ScZ vs. HC classification for RepOD dataset.

| Study | Method | Segmentation | Validation | Accuracy |
| --- | --- | --- | --- | --- |
| Shalbaf et al. [25] | ResNet-18-SVM | 5 | 10-fold | 98.60% |
| Shoeibi et al. [59] | CNN-LSTM | — | 10-fold | 99.25% |
| Das et al. [9] | SVM (Cubic) | — | — | 98.9% |
| Baygin et al. [26] | K-nearest neighbors (KNN) | — | 10-fold | 99.47% |
| Goker et al. [16] | 1D CNN | 5 min | — | 98.76% |
| Agarwal et al. [27] | Boosted trees classifier | 4 | — | 99.24% |
| Siuly et al. [24] | Deep Feature based SVM | — | 10-fold | 99.23% |
| Guo et al. [45] | Three-dimensional Convolutional Neural Network (3DCNN) | 4 s | 10-fold | 99.46% |
| Sahu et al. [37] | RDCGRU | 5 s | — | 88.88% |
| Lillo et al. [18] | CNN | — | — | 93% |
| Haider et al. [30] | PSDL | 225 segment per channel | — | 89.12% |
| Gosala et al. [31] | PSDL | 5s | 5-fold | 99.25% |
| Aziz et al. [4] | KNN | 20s | 10-fold | 99.4% |
| Buettner et al. [29] | RF | — | 10 | 96.77% |
| Jahmunah et al. [22] | SVM-RBF | 6250 point samples | 10-fold | 92.91% |
| De et al. [23] | SVM | — | — | 89.00% |
| Li et al. [32] | Microstate semantic model | — | 10-fold | 97.2% |
| Vazquez et al. [28] | RF | 644 segment per patient | 7-fold | 97.2% |
| Latreche et al. [34] | Ten-layer convolutional neural network(CNN) | — | 10-fold | 99.18% |
| **Proposed work** | **STFT based mel spectrogram CNN** | **3 s** | **10-fold** | **99.82%** |

“—” indicates not reported in the paper.

**Table 7.** Comparison with existing studies on ScZ vs. HC classification for kaggle basic Sensory ask dataset.

| Study | Method | Segmentation | Validation | Accuracy |
| --- | --- | --- | --- | --- |
| Sahu et al. [43] | Separable convolution attention network (SCZ-SCAN) via 2-D scalogram | 4 s | — | 95.00% |
| Siuly et al. [3] | EMD | — | 10-fold | 89.59% |
| Khare et al. [60] | RVMD and OELM | — | 10-fold | 92.93% |
| Khare et al. [38] | SPWVD-based TFR and CNN | 3s | 10-fold | 93.36% |
| Srinivasan et al. [61] | CNN+ LSTM | — | 10 fold | 98.2% |
| Guo et al. [21] | CNN | 3 s | 8:2 split | 92% |
| Ellis et al. [62] | 1D CNN+LSTM | 25 s | 10-fold | 75.9% |
| Ko et al. [17] | VGGNet- based CNN model | — | 10-fold | 93.2% |
| Barros et al. [8] | CNN | — | 10-fold | 78±8% |
| **Proposed work** | **STFT based mel spectrogram CNN** | **3 s** | **10-fold** | **98.31%** |

“—” indicates not reported in the paper.

Early studies often relied on conventional ML classifiers combined with handcrafted EEG features. For instance, Support Vector Machines (SVM) and k-nearest neighbors (KNN) proved unexpectedly competitive, achieving accuracies above 98% in several cases. Das et al. [9] demonstrated that an SVM model could achieve nearly 98.9%, while Baygin et al. [26] reported 99.47% with KNN. Similarly, boosted trees and random forests showed strong potential, with accuracies of 99.24% and 97.2%, respectively. These results indicate that even relatively simple classifiers can perform remarkably well when paired with effective feature engineering. Nevertheless, consistency across datasets was less reliable, as seen in other studies where tree-based methods and SVMs achieved closer to 89–92% accuracy. Shalbaf et al. [25] employed ResNet-18 to learn discriminative EEG features and then classified them using an SVM, reaching 98.6% accuracy. Siuly et al. [24] took a similar route, integrating deep feature extraction with an SVM classifier and achieving 99.23%. These findings highlight the value of leveraging the feature representation strengths of CNNs while preserving the stable generalization power of classic classifiers. Deep learning models—particularly CNNs and recurrent neural networks—have consistently achieved state-of-the-art results. Shoeibi et al. [59] used a hybrid CNN-LSTM framework and reported 99.25% accuracy. Guo et al. [45] advanced this further with a 3D CNN architecture designed, achieving 99.46%, one of the highest performances across the literature. Similarly, Latreche et al. [34] employed a ten-layer CNN that reached 99.18%. Beyond raw signal processing, spectral and time-frequency methods remain valuable for uncovering discriminative patterns. Power Spectral Density-based Learning (PSDL), for instance, showed mixed outcomes: Haider et al. [30] reported a modest 89.12%, whereas Gosala et al. [31] refined the same approach to achieve 99.25%.

Now, move on to the Kaggle dataset existing approaches, in Table 7, we can see that one notable trend is the use of time-frequency representations to capture the non-stationary characteristics of EEG signals better. For example, Sahu et al. [43] introduced a method that stood out by achieving a robust 95% validation accuracy.

Srinivasan et al. [61] demonstrated a high accuracy of 98.2% using 10-fold cross-validation. Siuly et al. [3] employed Empirical Mode Decomposition (EMD), achieving an accuracy nearing 90%. Building on this, Khare et al. [38] applied Recursive Variational Mode Decomposition (RVMD) combined with Orthogonal Extreme Learning Machines (OELM), pushing the accuracy higher to approximately 93%. Their subsequent work incorporated further boosting performance to over 93%. Parallel efforts by Guo et al. [21] also confirmed the utility of CNN models, reporting accuracy of 92%. Ellis et al. [62], for example, explored a hybrid 1D CNN and LSTM network with an accuracy of around 76%. Ko et al. [17] VGGNet-based CNN demonstrated reliable performance with approximately 93% accuracy. In comparison, Barros et al. [8] recorded lower accuracies around 78%.

Our proposed work in this line of research has taken time-frequency representation further by using STFT-based Mel spectrograms as inputs to CNNs. This approach achieved an outstanding 99.82% on the RepOD dataset and 98.31% for the Kaggle dataset, surpassing earlier frameworks and underscoring the strength of integrating signal decomposition with deep architectures. This trajectory suggests a future where clinically applicable EEG-based diagnostic tools for ScZ may be both feasible and reliable, and can be built through our system.

## Conclusion

In this research, we have developed a framework to identify critical brain lobes for the detection of ScZ from HC subjects using EEG data. Our approach begins by segmenting the EEG signals, followed by transforming these segments into mel-spectrogram images using STFT. These mel-spectrogram images are then classified using a CNN model. We evaluated our model on two datasets to bring clarity and acceptance of our proposed model. We applied this framework to the full brain lobe as well as five distinct brain lobes, aiming to pinpoint the regions most significant for ScZ detection. Through analyzing EEG data from publicly available two datasets, we have demonstrated a commendable accuracy in both datasets, where a mel-spectrogram image is used for the first time in this domain of research. We have also discovered that the frontal lobe showed the most significant changes across both the dataset, followed by the temporal lobes. The occipital lobe shows less importance in the diagnosis accuracy. These findings emphasize the importance of targeting specific brain lobes when developing diagnostic and monitoring tools for ScZ. To enhance the interpretability of our results and provide meaningful insights, LIME, SHAP, and Grad-CAM techniques is applied to improve the transparency of our framework further. By incorporating XAI techniques, our research not only advances the accuracy of ScZ detection but also provides actionable insights for clinicians and researchers to interpret the model’s decisions, ultimately contributing to more informed and effective healthcare solutions.

Research to date has largely focused on applying a single methodology across multiple diseases. Further studies should investigate the applicability of EEG in identifying and monitoring additional neurodegenerative disorders, including Parkinson’s Disease, Huntington’s Disease, and epilepsy, to evaluate the generalizability of our methodology. Therefore, there is an opportunity to extend and validate this approach by implementing it in the context of other disorders. Integrating multimodal data, including structural MRI and genetic information, could boost the model’s predictive capability and provide a more thorough explanation of disease causes.

Although we have already evaluated our technique on two distinct datasets, testing it on additional large EEG datasets for ScZ will enhance its generalization. Ultimately, the incorporation of EEG-based frameworks into clinical practice represents a potential breakthrough. Executing our methodology in practical environments and performing longitudinal research could result in critical insights, facilitating the enhancement of the technology and ultimately advancing personalized, effective healthcare solutions for neurological diseases.

## Author Contributions

**Conceptualization**: Md. Milon Hossain, Md. Nurul Ahad Tawhid.

**Data curation**: Md. Milon Hossain.

**Formal analysis**: Md. Milon Hossain, Md. Nurul Ahad Tawhid.

**Investigation**: Md. Milon Hossain.

**Methodology**: Md. Milon Hossain, Md. Nurul Ahad Tawhid.

**Resources**: Md. Milon Hossain, Md. Nurul Ahad Tawhid.

**Supervision**: Md. Nurul Ahad Tawhid.

**Visualization**: Md. Milon Hossain.

**Writing – original draft**: Md. Milon Hossain

**Writing – review & editing**: Md. Milon Hossain, Md. Nurul Ahad Tawhid

## Notes

### Competing Interest Statement

The authors have declared no competing interest.

